# DeepPathway: Predicting Pathway Expression from Histopathology Images

**DOI:** 10.1101/2025.07.21.665956

**Authors:** Muhammad Ahtazaz Ahsan, Karen Piper Hanley, Martin Fergie, Claire O’leary, Gerben Borst, Federico Roncaroli, Fayyaz Minhas, Magnus Ratrray, Mudassar Iqbal, Syed Murtuza Baker

**Affiliations:** Division of Informatics, Imaging and Data Sciences, Faculty of Biology, Medicine and Health, The University of Manchester, UK; Division of Diabetes, Endocrinology & Gastroenterology, Faculty of Medical Sciences, The University of Manchester, UK; Division of Cancer Sciences, Faculty of Biology, Medicine and Health, The University of Manchester, UK; Division of Neuroscience, Faculty of Biology, Medicine and Health, The University of Manchester, UK; Department of Computer Science, University of Warwick, Coventry, UK

## Abstract

Spatial transcriptomics (ST) technologies provide spatially resolved gene expression along with image data, allowing the integrative analysis of complex tissue microenvironments. Despite their potential, the widespread adoption of ST remains limited due to high costs, and methodological challenges in data acquisition. Thus, there have been recent efforts to develop deep learning methods capable of inferring spatial gene expression from the much cheaper and easily available haematoxylin and eosin (H&E) images. These methods demonstrate promising results in reconstructing transcriptomic landscapes within tissue sections. While existing approaches predominantly focus on gene-level predictions, biological processes are often regulated at the pathway level through coordinated activity among functionally related genes. We present DeepPathway, a contrastive learning-based approach trained on ST data to predict pathway expression from H&E-stained sections. We compute input pathway expression by summarizing the expression of constituent genes using established pathway definitions. We evaluate the performance of our method on two prostate cancer datasets and validate our approach on the H&E images acquired from The Cancer Genome Atlas (TCGA) clearly differentiating between normal and tumour tissues. Finally, we apply our method to predict hypoxia signatures using H&Es of brain tumour samples where hypoxia staining with pimonidazole was available as ground truth. Implementation code for DeepPathway is available at https://github.com/aahsan045/DeepPathway.

## I. Introduction

Tissues in multi-cellular organisms consist of diverse cell populations that coordinate their functions by spatially interacting with each other and with the microenvironment [1]. Single-cell RNA sequencing (scRNAseq) technologies allow high-throughput gene expression profiling to study individual cellular activity but lack the spatial context. To address this limitation, spatial transcriptomics (ST) [2]) technologies, such as the Visium platform by 10x Genomics and Slide-seq [3] have been developed. These methods retain the spatial context of cells while capturing gene expression data without disrupting the complexity of tissue organisation. ST platforms often include co-registered Haematoxylin and Eosin (H&E)-stained images providing a view of tissue architecture and heterogeneity at microscopic level. The integration of pathological, ST and molecular data enables researchers to accurately analyse tissue regions that can be relevant to diagnosis, disease progression or are driven by molecular pathways that can be targeted with effective treatment modalities [4].

Although ST provides detailed molecular insights, it is an expensive and time-consuming technique that requires unique expertise. In contrast, tissue scanned H&E-stained images are cheap and readily available, from routine diagnostic provision. Recent advances in deep learning (DL)-based methods have enabled the prediction of spatial gene expression directly from histology images [5]–[19] These models are trained on ST datasets and then used to infer gene expression patterns from H&E images alone. For example, ST-Net [5] utilises DenseNet-121 (pretrained on ImageNet) for extracting features from H&E images and predicts expression levels for 250 genes. Similarly, BLEEP (Bimodal Embedding for Expression Prediction), proposed by Xie et al. [6], employs a bimodal contrastive learning framework [20] to learn a shared embedding space between H&E patches and corresponding gene expression profiles. Despite being trained on thousands of genes, these methods tend to predict only a limited subset of disease-relevant genes with reasonable accuracy.

Normal cell functions result from the coordination of complex, hierarchical gene networks^1^. The analysis of such networks and pathways rather than individual genes reflects more closely the tissue homeostasis in normal and pathological conditions. To translate a set of individual gene expression profiles into a pathway expression profile at cellular or spot level resolution, methods such as AUCell [22] or Ucell [21] can be utilised. This provides a way to transform thousands of genes into a handful of relevant pathways which could be more accurately and efficiently learned and could have more translational impact in identifying biomarkers for diagnosing certain disease phenotypes.

With the aim to predict the pathway expression from H&E images, we develop a DL-based approach called DeepPathway, adapted from BLEEP [6]. BLEEP is a contrastive learning framework that aims to learn a joint embedding space of expression and image modality via contrastive loss. However, BLEEP has not been developed for pathway expression prediction problem. Also, it requires access to ST training data (or its embeddings) to infer the expression of test histology image, somewhat limiting its practical deployment. Hence, we introduce a novel two-stage model for pathway expression prediction replacing BLEEP’s inference mechanism with a deep neural network (DNN) that directly predicts pathway expression from H&E features. The key contributions of this work are given below.

1. We demonstrate the effectiveness of pathway expression prediction over gene expression prediction using prostate cancer datasets. For this we introduce a new approach adapted from an existing bimodal contrastive learning framework to predict the pathway expression from H&E images.
2. We propose a two-stage learning framework, DeepPathway, to predict pathway expression directly from H&E image.
3. We validate our approach on a larger prostate carcinoma ST dataset. Additionally, we show the application of the proposed method on the H&Es of the prostate tumour samples from The Cancer Genome Atlas (TCGA).
4. Finally, we use the proposed method to identify hypoxic regions in brain tumour patient samples using only H&E images which were validated using hypoxia specific staining with pimonidazole.

## II. Materials and Methods

### A. Approach overview

A brief overview of DeepPathway is presented in Fig. 1 which shows as input paired H&E images and gene expression data coming from ST experiments. After initial gene expression and H&E image processing, using literature-based pathway definitions (e.g., MSigDB hallmark), we transformed the original spot-by-gene matrix into a spot-by-pathway matrix using UCell [21]. We train DeepPathway on paired image patch and pathway expression data to learn a common embedding representation via contrastive learning. Finally, we use trained DeepPathway to predict the pathway expression of test H&E images.

**Fig. 1:**
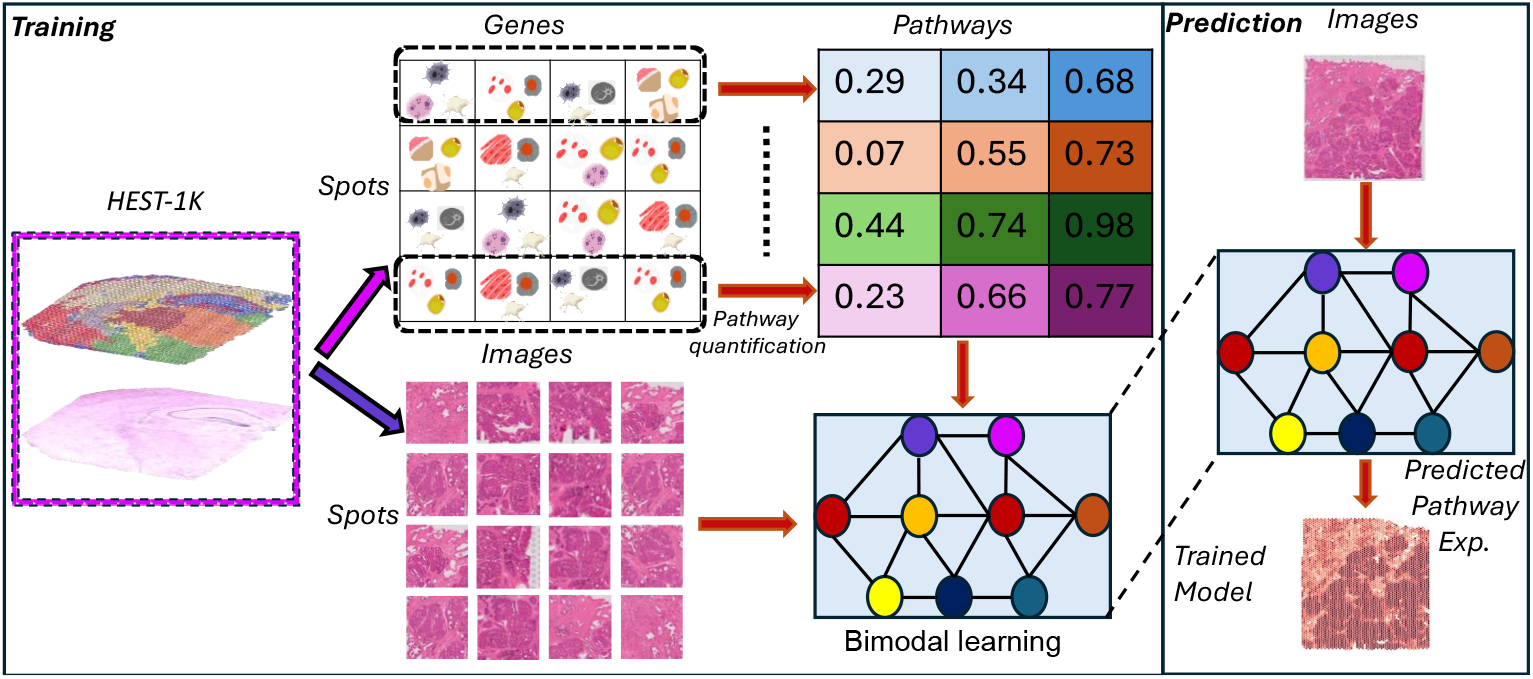
Overview of the proposed method (**DeepPathway**); Spot-level gene expression and H&E image patches are acquired from HEST-1k database. Gene expression data is summarized into spots × pathway expression matrix using pathway quantification with UCell [21]. In the training part, a deep learning model is trained using bimodal contrastive learning on spot-level image patches and pathway expression data. At the inference time, only H&E image is passed to the trained model for predicting pathway expression.

We organised the work reported here into three interconnected approaches, which are briefly described as follows. (1) At first, we used BLEEP as the baseline method to perform and compare gene expression prediction followed by conversion into pathway expression versus direct pathway expression prediction highlighting the utility of our approach. (2) In the second approach, we improved image feature representations by integrating outputs from a trainable BLEEP image encoder with features from the non-trainable H&E image foundation model, *H-optimus-0* [23]. Features extracted from *H-optimus-0* foundation model help ResNet18 learn more meaningful image representation when used as prior knowledge. (3) Finally, we replace the inference part of BLEEP with training a deep neural network (DNN) model for improved deployability. This results in a proposed two-stage learning framework in which, first a model having an image encoder with optimus-and expression encoder is trained using contrastive learning (stage-1). In stage-2, we train a DNN model on image features extracted from the image models used in the stage-1 to predict the pathway expression directly(see Fig. 3A).

Throughout all stages, we leveraged the bimodal contrastive learning framework introduced in BLEEP [6] that was originally developed for predicting spatial gene expression from H&E images. The model takes randomly shuffled batch of samples for training. The main objective is to learn the same-sized embedding space between matching image-expression pairs. As model training continues, the model improves its performance by capturing meaningful cross-modal similarities while pushing apart dissimilar pairs, thereby enhancing the modality alignment. In the original BLEEP inference setup, test image embeddings are compared to those from the training set, and the pathway expression is estimated by averaging the outputs from the top k closest matches. In contrast, our two-stage framework eliminated the need for training data (or embeddings) at inference time and pathway expressions are predicted directly from the image features using the trained DNN.

### B. Bimodal contrastive learning for pathway expression prediction

Contrastive learning (CL) [24], [25] is a computational method applied to learn meaningful semantic representation by pulling similar samples closer to each other while pushing dissimilar samples apart. This idea has been extended to bimodal data such as image-text, to learn a joint feature representation across modalities. Additionally, it is widely used for developing DL-based applications such as image-caption generation [26] or text-to-image generation [27]. Adapting the idea of CL from BLEEP, given a spot (a spatially barcoded area where gene expression is profiled) *S* = {*X*_*img*_, *X*_*expr*_}, that consisted of paired image-expression profile, a learnable image encoder (*f*_*θ*_(*X*_*img*_)) and an expression encoder (*f*_*ϕ*_(*X*_*expr*_)) are used to produce embeddings *Z*_*img*_ and *Z*_*expr*_, respectively. We then add the positional encoding of the corresponding spot location coordinates (*x, y*) into both embeddings by adapting idea of sinusoidal positional encoding (*f*_*PE*_ (x,y)) as proposed for transformers [28]. Finally, we calculate the contrastive loss using Equation 1.

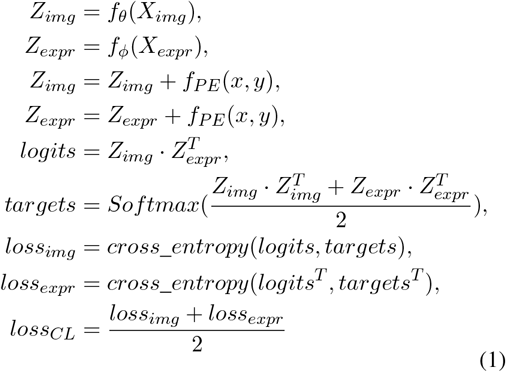

where *logits* is the paired similarity vector between both modalities treated as a ground truth and *targets* is the averaged internal similarity between both modalities followed by *softmax*. Then cross-entropy is applied between them to calculate the image and expression loss followed by taking their mean for producing contrastive loss. The final loss (Ł) is calculated by adding *λ* scaled *L*_1_ regularization to both image and expression embeddings,

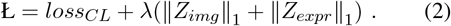

### C. Dataset

We obtained publicly available 10X Visium datasets from the HEST-1k database [29], which provides uniformly processed high quality images, gene expression profile, metadata for each ST sample, covering multiple organs and diseases. Specifically in HEST-1k, all whole-slide images (WSIs) are normalized and transformed to generic Pyramidal-based TIFF image objects that can be integrated with computational tools such as OpenSlide [30] or image viewers like QUPath [31]. In this work, we used two different human prostate cancer datasets from HEST-1K, one we refer to as small prostate cancer dataset [29] and other as large prostate cancer dataset [32] hereafter. The large data from prostate adenocarcinoma consisted of two patients, patient 1 has 8 samples while patient 2 has 15 specimens, all diagnosed as Gleason 4+3 score group 3. Additionally, we used skin cancer dataset [33], consisted of 4 ST samples. or prediction of hypoxia pathway, we used 5 glioblastoma multiforme (GBM) cancer ST samples from Ravi et.al [34]. Details about total number of samples used for each dataset, total number of spots left after preprocessing, and image resolution of whole slide images (WSIs) for each dataset can be seen in Table I.

**TABLE I:** Statistics for the datasets used in this study.

| Sr. No. | Dataset | Number of Samples | Total Number of Spots | Image Resolution ($\mu\text{m}/\text{pixel}$) |
| --- | --- | --- | --- | --- |
| 1 | Small prostate cancer | 5 | 19245 | 0.61 |
| 2 | Large prostate cancer | 8 (Patient 1)<br>15 (Patient 2) | 18965 (Patient 1)<br>34845 (Patient 2) | 0.57, 0.68, 0.17 (Patient 1)<br>0.34 (Patient 2) |
| 3 | Skin cancer | 4 | 8839 | 0.17 |
| 4 | Brain tumour (GBM) | 5 | 17425 | 0.46 |

### D. Data processing

For each dataset, we performed gene filtering to remove genes that are expressed in less than 10 spots and spot filtering to remove spots that have less than 500 genes expressed. Then we selected the top 10,000 highly variable genes (HVG) from each sample. The reason for selecting a large number of HVGs is to keep the maximum overlap between each sample’s genes with the pathway-specific genes. For pathways definitions, we used the Molecular Signature Database (MSigDB) hallmark collection [35], which is commonly used for cancer studies. It consists of 50 pathway definitions with the unique number of genes equal to 4383. After acquiring the spot-level gene count data, we performed the total normalization followed by the *log1p* transformation using *scanpy* [36]. Normalizing the data deals with different sequencing depths and helps in comparison across spots while the *log1p* transformation reduces the skewness in the data.

Then we performed the H&E image processing on the spots left after gene expression processing. For each dataset, we cropped the image patch of 224 *×* 224 pixels centered at each spot’s location using the given resolution of microns per pixel through the TIA toolbox [37]. We removed spots where more than 75% of the image represented the background, i.e., having pixel value in all three color channels more than 200. For GBM samples from Ravi et al. [34], we applied Reinhard H&E image stain normalization to obtain the same histology colors distribution and for smooth model training. For GBM and TCGA H&E image data only, we processed WSIs as follow. Starting from the at top-left corner, i.e., at (0,0) location of the image, non-overlapping patches of 224 × 224 were taken and passed to the image processing pipeline to filter out non-informative patches (as mentioned above). In case of GBM data, informative patches were then Reinhard normalized as training data consisted of different colored H&E images. During inference, same image processing is applied to predict pathway expression while their locations were preserved for visualization. Additionally, we have integrated the SpatialData [38] in DeepPathway’s code, which provides a Python-based implementation of handling spatial omics data data, for unified data storage and visualization.

### E. Pathway expression quantification

We used UCell [21] to quantify the pathway expression given the gene expression measurements and pathway definitions from MSigDB hallmark. UCell is a recent pathway quantification method which is time and memory efficient compared to AUCell. It uses Mann-Whitney statistics to compare the rank distribution of genes in a pathway versus all other genes. It is robust to differences in dataset compositions as it only considers relative gene expression in a given spot/cell and avoids an arbitrary threshold which is the case in AUCell. In this method, genes in each spot are ranked (from high to low) based on their expression values. Then, for a *j*^*th*^ spot, the UCell score (*U*_*j*_) for a pathway composed of *n* genes is calculated using Eq. 3.

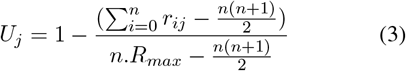

where *r*_*ij*_ is the rank value of *i*^*th*^ the gene in the *j*^*th*^ spot. The *R*_*max*_ is the maximum rank value used to drop the uninformative tail of genes with low expression. We selected the *R*_*max*_ value for each dataset independently using the following method. For a given spot within a sample, we sorted the gene expression from highest to lowest values and obtained the rank index of the first zero value. We repeated this for all spots across all samples and took the median to obtain the final *R*_*max*_ threshold. The reason for this approach to select *R*_*max*_ threshold is to give constant rank value for all non-expressed genes to mitigate the long tail of bottom ranking genes. By following this procedure, we found the *R*_*max*_ value of 4276, 1926, 1954, 1265, and 2106 for small prostate, large prostate cancer patient 1, large prostate cancer patient 2, skin cancer, and GBM cancer data, respectively. Also, we selected only those pathways that contained at least 70% of the genes overlapping with the ST expression dataset genes (that are union of all samples’ genes in the dataset) and contain minimum 20 genes. By using this threshold, we found the number of pathways for small, large prostate cancer, and skin cancer datasets to be 50, 47, and 44, respectively. For GBM data, we used 7 brain tumor specific pathway definitions.

### F. Model training and validation

Using a dataset of ST samples consisting of H&E image patches and their corresponding expression profiles, we assessed the performance of each model using leave-one-out cross validation, where one sample from the data is held out for testing while remaining data (comprising multiple tissue samples) is divided into spot-level training and validation data with a split ratio of 80% and 20%, respectively. We used ResNet50 for gene expression prediction experiments as it gives 2048-dimensional feature vector to train and predict thousands of genes while we used ResNet18 (which provides 512-dimensional feature representation) for pathway expression prediction due to its lower computational complexity as well as suitable choice to train and predict a few pathways. For gene expression prediction task, we used the union of top 10,000 highly variable genes from each sample for each dataset. Models were trained using the Adam optimizer with a learning rate of 0.001 and training batch size of 256. We kept the same embedding size of 256 for input image projection, gene/pathway expression projection, and positional encoding data. We also integrated pre-trained *H-optimus-0* (a vision-transformer model, trained on more than 500, 000 H&E stained whole slide histology images), in our experiments which provides an embedding of 1538-dimensional vector for each image patch. All performance metrics are evaluated on test data. For pathway expression prediction experiments, we used the criterion of early stopping with a patience level of 5 to monitor the improvements in validation loss to prevent overfitting. We used the value of *λ* = 0.0001 to control the strength of *L*_1_ penalty to encourage sparsity in both image and expression embeddings while reducing noise. To access the quality of predicted expression, we used the Pearson correlation coefficient (PCC), as well as spatial visualization of ground truth and predicted expression.

## III. Results

### A. Bimodal contrastive learning for gene versus pathway expression prediction on prostate cancer data

To being with, we applied BLEEP to perform gene expression prediction followed by conversion to pathway expression and direct pathway expression prediction on the small prostate carcinoma dataset to establish the utility of pathway-level prediction while using the same underlying contrastive learning approach. At first, spot-level paired gene and H&E image data is processed and BLEEP having ResNet50 as an image encoder is trained on paired data to predict gene expression. This trained model is then used to predict gene expression for test sample, which is then converted into pathway expression by Ucell, utilising MSigDB hallmark pathway definitions (we call this approach “BLEEP-genes”). In parallel, we passed the processed gene expression data along with the MSigDB pathway definitions to UCell to obtain pathway expression data. Then, we trained BLEEP using same ResNet50 model (image encoder) with paired pathway expression data to predict the pathway expression (we call this approach “BLEEP-pathways”). We used ResNet50 only here in BLEEP-pathways to make a fair comparison with BLEEP-genes. Finally, reconstructed pathway expression obtained from BLEEP-genes and predicted pathway expression from BLEEP-pathways are compared (Fig. 2A) by computing the PCC between the ground truth pathway expression of the test sample—and both the reconstructed and predicted pathway expressions. Overall, BLEEP-pathways clearly outperformed BLEEP-genes (Fig. 2B). For example, in the case of Androgen Response (Fig. 2C), BLEEP-pathways has PCC of 0.63 while BLEEP-genes with 0.13. On the other hand for some pathways, BLEEP-pathways and BLEEP-genes performed similarly, for example Epithelial Mesenchymal Transition (EMT) pathway, which plays a crucial role in tumour metastasis [39], for samples “INT27” and “INT28” have PCC values of around 0.79 and 0.55, for both the approaches, respectively. This potentially depicts that genes in the EMT pathway are well correlated with the histological microscopic features, helping predicting the genes and its constituent pathway very well in both scenarios. Additionally, we computed the PCC between top 5 highly expressed and highly variable pathways from BLEEP-genes and BLEEP-pathways (Supplementary Fig. 1). It can be seen that BLEEP-pathways is consistent with higher PCC values across all samples, which depict that it is more meaningful to perform direct pathway expression prediction as it leverages the advantage of focusing on aggregate biological signal rather than individual gene fluctuations. Furthermore, for the sake of a fair and thorough comparison, we evaluated BLEEP-genes and BLEEP-pathways on ResNet18 as an image encoder (Supplementary Fig. 2), showing similar results of BLEEP-pathways performing overall better in most of the samples compared to BLEEP-genes.

**Fig. 2:**
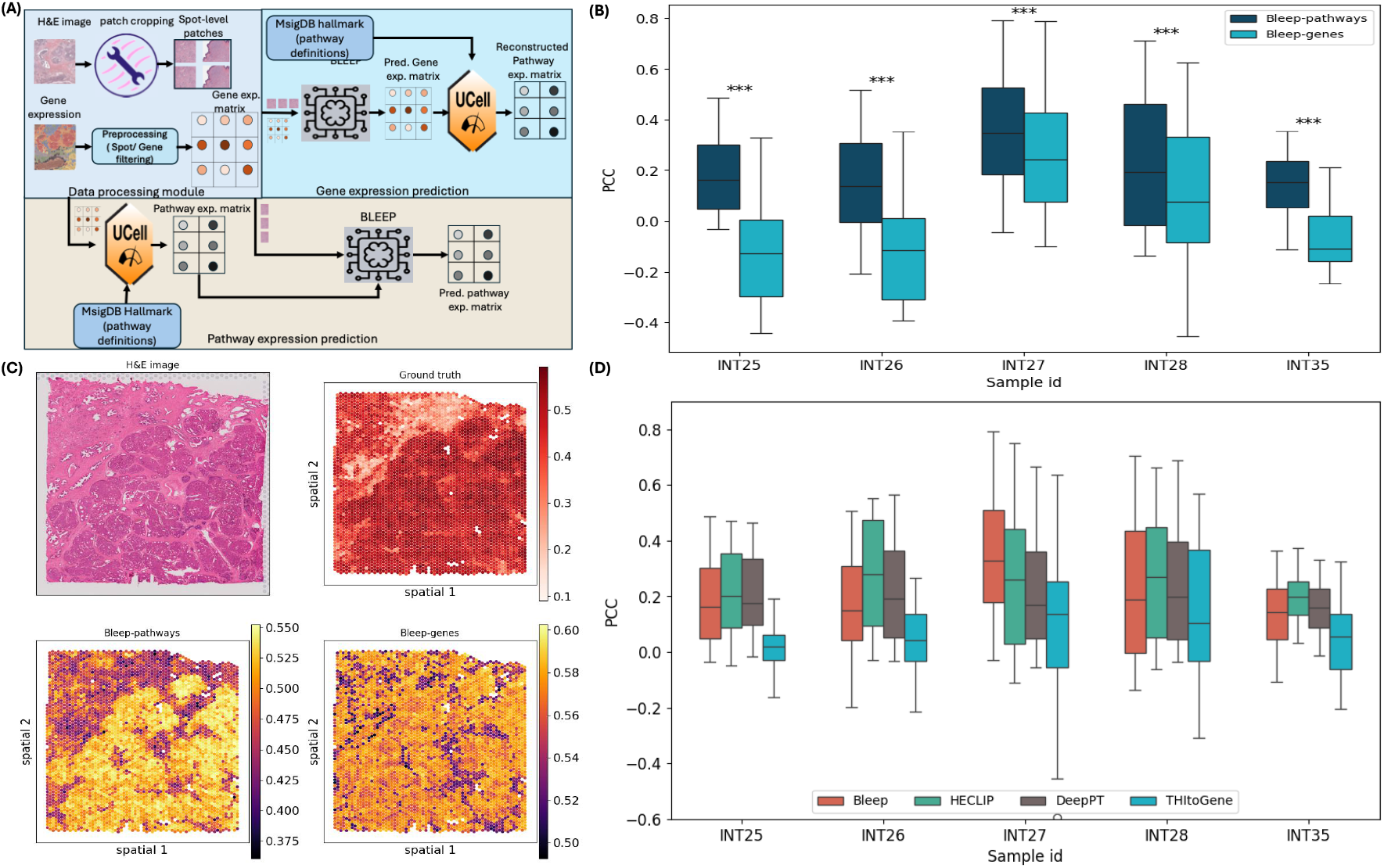
**(A)** Overview of the baseline method where spot-level data is initially processed (data processing module). Then, BLEEP is first trained on paired H&E images and gene expression data to predict the gene expression. Then predicted gene expression is passed to the UCell for obtaining the pathway expression. Whereas in pathway expression prediction (bottom half), gene expression matrix is first converted into pathway expression matrix followed by training BLEEP to predict the pathway expression. Finally, the pathway expression matrix from gene expression prediction (BLEEP-genes) is compared with the predicted pathway expression (BLEEP-pathways) obtained from pathway expression prediction. **(B)** PCC distribution plot of BLEEP-pathways versus BLEEP-genes for all samples where “*” symbol shows the magnitude of p-value calculated from paired t-test of two PCC distributions (“***” for p-value *<*0.001). **(C)** Spatial visualization of ground truth pathway expression from UCell, predicted *Androgen response* pathway expression from BLEEP-pathways and BLEEP-genes having PCC value of 0.63 and 0.13, respectively, with H&E image (**INT28**). **(D**) PCC distribution plot showing comparison of BLEEP with other existing methods in predicting pathway expression. All experiments were conducted on small prostate cancer dataset.

**Fig. 3:**
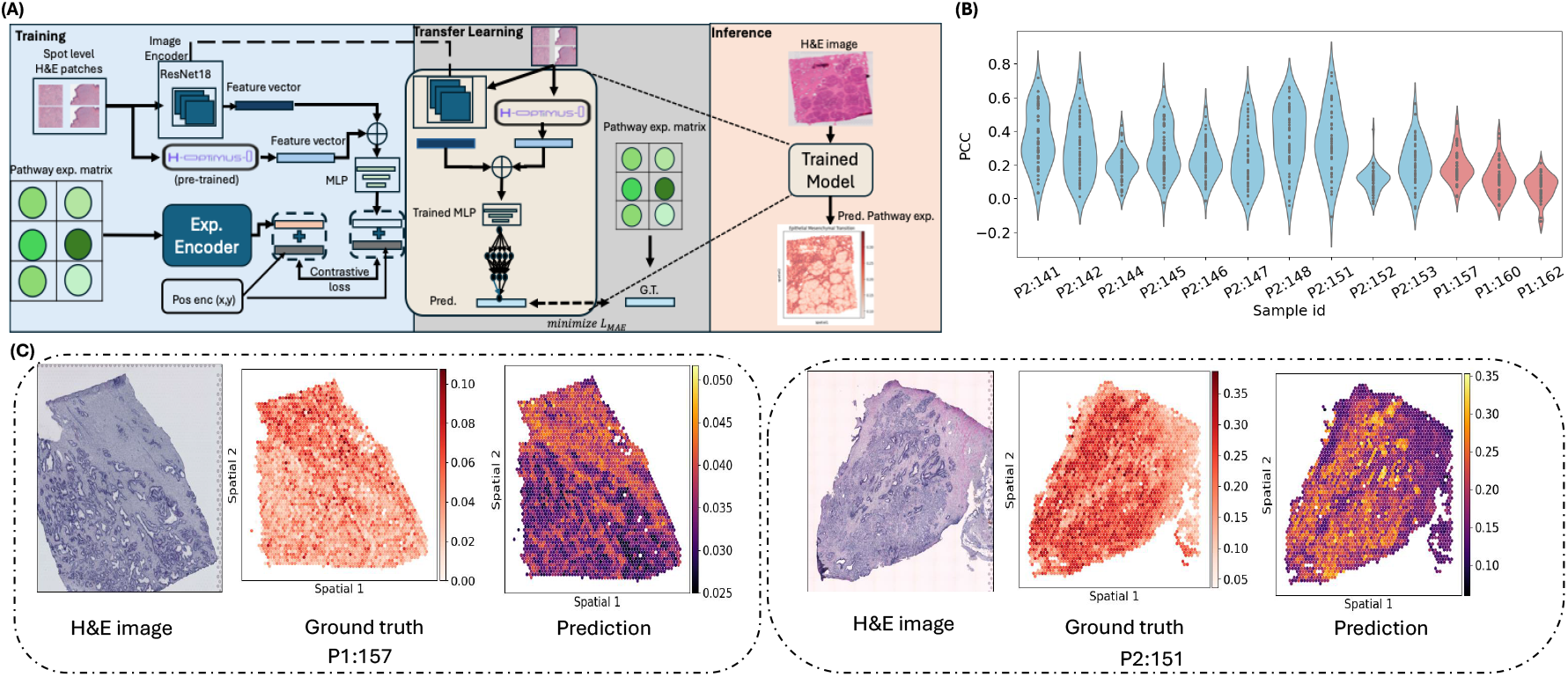
**(A)** Overview of DeepPathway: the proposed two-stage learning framework. At first, ResNet18 is trained on combined image features that are obtained from ResNet18 and *H-optimus-0* foundation model using bimodal contrastive learning. In second stage, a DNN is trained on image features extracted from trained stage-1 model to predict pathway expression by minimizing MAE. During inference time, only H&E image is used to predict pathway expression. **(B)** PCC distribution plot of test samples from patient 1 (P1) and patient 2 (P2) when the model is trained using the training samples from both patients. **(C)** test H&E image and spatial visualization of the IL-6/JAK/STAT3 Signaling pathway from P1 and Androgen Response pathway from P2, shows the ground truth pathway expression—obtained from UCell—alongside the predicted pathway expression from the trained model. The predicted expression achieved PCC values of 0.46 and 0.75, on P1:157 and P2:151 samples, respectively.

In order to select the most suitable gene expression prediction model for adaptation to pathway expression prediction as well as using it as baseline for further improvements, we performed comparison of BLEEP with existing methods, i.e., HECLIP [7], DeepPT [40], and THItoGene [10]. DeepPT and THItoGene are selected from top performing gene expression prediction methods, as reported in the benchmarking study [41]. These were adaptable to pathway expression prediction task given their underlying model architectures while others like Hist2ST [12] used zero-inflated negative binomial loss for the gene expression prediction task, which is not suitable for pathway expression prediction. We also included recently published method HECLIP in the comparison as its architecture is very similar to BLEEP model. We evaluated each model on small prostate carcinoma data and the PCC distribution of each test sample is shown in Fig. 2D. Our results show that BLEEP outperformed other methods while achieved similar performance as HECLIP. Thus, we preferred BLEEP over other methods and built our proposed model (explained in next section) using BLEEP as a baseline model.

### B. Improved pathway expression prediction model deployability via two-stage learning framework

One of the limitations of the BLEEP model is that it requires the training data (latent projections) to infer the predicted pathway expression for the test data. BLEEP predicts expression by projecting test image into the embedding space and retrieving the average of top k matched candidates from the reference (training) dataset. However, this might not always be feasible considering future deployability of these models in translational settings. Thus, we propose a two-stage learning framework (Fig. 3A) where we first trained ResNet18 as an image encoder with *H-optimus-0* using contrastive learning (stage-1) followed by training a DNN model by giving these features obtained from stage-1 model as an input to predict a vector of pathway expression. This approach minimizes the mean absolute error (MAE) between the ground truth and prediction, which is calculated as 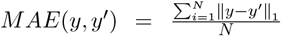, where *y* is ground truth pathway expression obtained from UCell and *y*^*′*^ is the predicted pathway expression for a batch of *N* samples. Additionally, here we did not add the positional encodings, as we are obtaining the image features from the trained model which has already learned the spatial similarities between those modalities. During inference, we used the trained ResNet18, *H-optimus-0* (stage 1), and the trained DNN model (stage 2) to predict the patch-level pathway expression. Beside improved deployability, adding a DNN in the two-stage architecture makes the model lightweight for prediction and open ups the way of developing transfer learning based applications in future. We did an ablation study evaluating different combinations of the proposed two-stage model, i.e., BLEEP with Optimus, BLEEP with DNN and ResNet18 with Optimus and DNN (which is our final model and we call it as DeepPathway) on small prostate cancer data where DeepPthway was the best performing among all (Supplementary Fig. 3). Subsequently, we applied DeepPathway on large human prostate carcinoma data, classified as Gleason 4+3, grade group 3 [32] using two approaches, which are described as follows. Within the large human prostate data, we picked two patients with largest number of samples and used 5 random samples from each (a total of 10 samples) as training data for DeepPathway, and then performed testing on remaining 13 samples of both patients. Each patient was processed independently using their own ST data for fair comparison. In second approach, we used only patient specific data using leave-one-out cross validation. We evaluated model’s performance stage-wise, i.e.,stage-1 followed by stage-2 using both approaches separately to show the model’s robustness and generalisation. Results for both approaches are described below.

#### 1) Evaluation on mixed patient’s samples

The results for stage-1 model using mixed patient samples are shown in Supplementary Fig. 4 where variation in PCC distributions depicts differences in model performance across different patient samples, potentially pointing to intrinsic heterogeneity within the tissue which could influence model’s performance. Additionally, we evaluated stage-1 approach on small prostate cancer data (Supplementary Fig. 5). These results show that our proposed approach achieved improved results in terms of pathway expression prediction with the inclusion of the *H-optimus-0* foundation model.

For stage-2, i.e., training a DNN for direct prediction with mixed patient data, PCC distribution between ground truth and predicted pathway expression is shown in Fig. 3B. A similar trend of PCC distribution is observed when compared to Supplementary Fig. 4B, as the bimodal contrastive loss model remained same in both approaches. Two different pathways (selected from the top 5 highly predicted), i.e., IL-6/JAK/STAT3 Signaling pathway from P1 (ID=157) and Androgen Response pathway from P2 (ID=151) with PCC values of 0.46 and 0.75 respectively, are visualised in Fig. 3C. This shows that our proposed approach not only aligns the image-expression modality using contrastive learning but also learned a meaningful association of image features with pathway expression through a simple DNN model.

#### 2) Evaluation using patient specific data with leave-one-out cross validation

Here, DeepPathway model’s stage-1 (Bleep with Optimus) is evaluated using patient-specific leave-one-out approach (Supplementary Fig. 6) which shows that for P1, few pathways have achieved PCC values of up to 0.40 even when the ground truth in that patient has very low spatially localized patterns, while, for P2, some samples have achieved high PCC values, e.g. 0.83 for EMT, 0.81 for Myogenesis, and 0.60 for Androgen Response. Similarly, for stage-2 model, results are shown in Fig. 4 which shows similar trend as observed in case of stage-1, i.e., the model has performed reasonably well in predicting pathway expression on most of the test samples of P2, while on P1 samples, only few pathways are predicted well with reasonable PCCs. For example, Myogenesis pathway is predicted with the PCC of 0.47 (ID=158), whereas the same pathway is predicted with the PCC of 0.77 in P2 (ID=150). This behavior shows that the model aligns well in a bimodal setting where the spatially localized patterns are present in both H&E and pathway expression. To support this further, we computed the inter-pathway correlation (PCC between each pathway with other pathway across all spots) using the predicted pathway expression obtained from DeepPathway, heatmap for the best predicted samples having ID=158 from P1 and ID=150 from P2, are shown in Supplementary Fig. 8 and Supplementary Fig. 9, respectively. It can be seen that in case of P1, there is no pathway-expression level heterogeneity, which can be learned with histopathological features in a contrastive learning manner. Almost, all pathways in ground truth show either a PCC of around +0.25 and −0.25 with each other, making it harder for model to learn (in mixed sample approach as well as in leave-one-out approach). However, for P2, there is pathway expression level heterogeneity. For instance, for both ground truth and prediction (as shown in Supplementary Fig. 9), p53 pathway and mTORC1 Signaling pathway are positively correlated. The p53 is a tumour suppressor pathway and maintains the cell integrity by activating cell cycle arrest, DNA repair, or apoptosis in case of DNA damage. The mTORC1 is the regulatory pathway for cell growth and metabolism. Thus, in Gleason 4+3 prostate cancers, both pathways can co-exist to promote tumour growth as they contribute in cell survival and proliferation. Furthermore, EMT and Myogenesis pathways are also positively correlated in both ground truth and predictions, co-exist in prostate cancers and contribute to tumour diversity, specifically, with respect to metastatic potential and differentiation state.

**Fig. 4:**
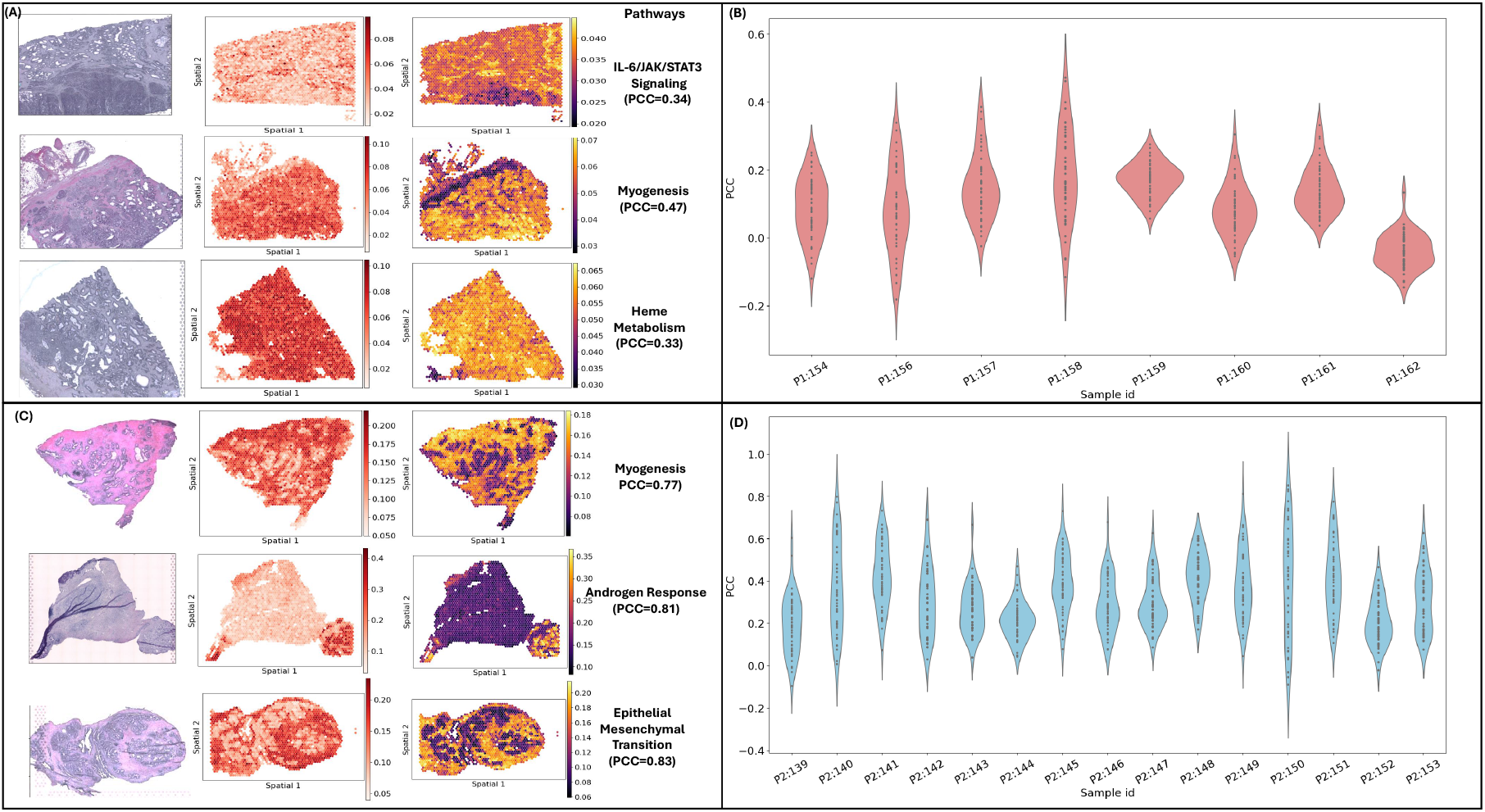
**(A)** Spatial Visualization of selected pathways (from top 5 predicted pathways) when DeepPathway model is evaluated using leave-one-out cross validation using samples of Patient 1 (P1) only. **(B)** PCC distribution of each test sample in P1 data when DeepPathway model is trained only on P1 samples. **(C)** Spatial Visualization of selected pathways (from top 5 predicted pathways) when DeepPathway model is evaluated using leave-one-out cross validation using samples of Patient 2 (P2) only. **(D)** PCC distribution of each test sample in P2 data when DeepPathway model is trained only on P2 samples.

We also analysed the pathways predicted well within each tissue sample individually using both approaches. These pathways are selected from top 5 highly predicted pathways based on PCC scores. Spatial visualization of test samples evaluated from mixed sample DeepPathway model, leave-one-out approach of P1 and P2, are shown in Supplementary Fig. 10, Supplementary Fig. 11, and Supplementary Fig. 12, respectively. Additionally, we evaluated stage-1 and stage-2 of DeepPathway model on skin cancer dataset (Supplementary Fig.7 A-C), where 4 ST samples are evaluated using leave-one-out cross validation.

For the sake of evaluating DeepPathway on external and independent datasets, we used two prostate cancer ST samples [42] from HEST-1k from a different study to predict the pathway expression. This dataset is processed independently, and we found out 20 out of 50 pathways present in the data. We predicted pathway expression from both models, i.e., using mixed patient samples model and best predicted model of P2 in leave-one-out settingd We used mixed patient samples model and the best model (having sample ID=150) and predicted 47 pathway expression. Then we sliced those 20 pathways from predicted expression which are available in ground truth for spatial visualization and PCC computation (see Supplementary Fig. 13). Additionally, we used small prostate cancer dataset as a part of external validation (see Supplementary Fig. 14).

### C. Pathway expression mapping in prostate cancer using TCGA whole slide images

We extended the application of our model by predicting pathway expression from prostate cancer dataset in the cancer genome atlas (TCGA); the most widely used public cancer data resource. We acquired WSIs of 8 patients which have both normal and prostate tumour tissue slides from TCGA data portal. We then used the best performing weights of our proposed model to predict the patch-wise pathway expression using only WSIs. We selected three representative pathways, i.e, EMT, Monogenesis, and Androgen Response, for patch-wise visualisation. EMT drives tumour metastasis [39] while activation of androgen receptor signalling drives proliferation and survival of prostate cancer cells [43]. Finally, stromal changes in prostate carcinoma including the activation of the myogenesis programme in smooth muscle cells are now recognised to correlate with poor outcomes [44]. Fig. 5 shows the results of the predictions obtained for these three pathways on two TCGA tumour patients (row-wise). The results for remaining tumour patients and their normal tissue sections are shown in Supplementary Fig. 15 and Supplementary Fig. 16, respectively. Myogenesis and Androgen Response show an inverse behavior while EMT and Myogenesis have similar behaviour with respect to their expression values in both normal or tumour samples. However, the expression intensity as well distribution of each pathway in normal section remains almost same.

**Fig. 5:**
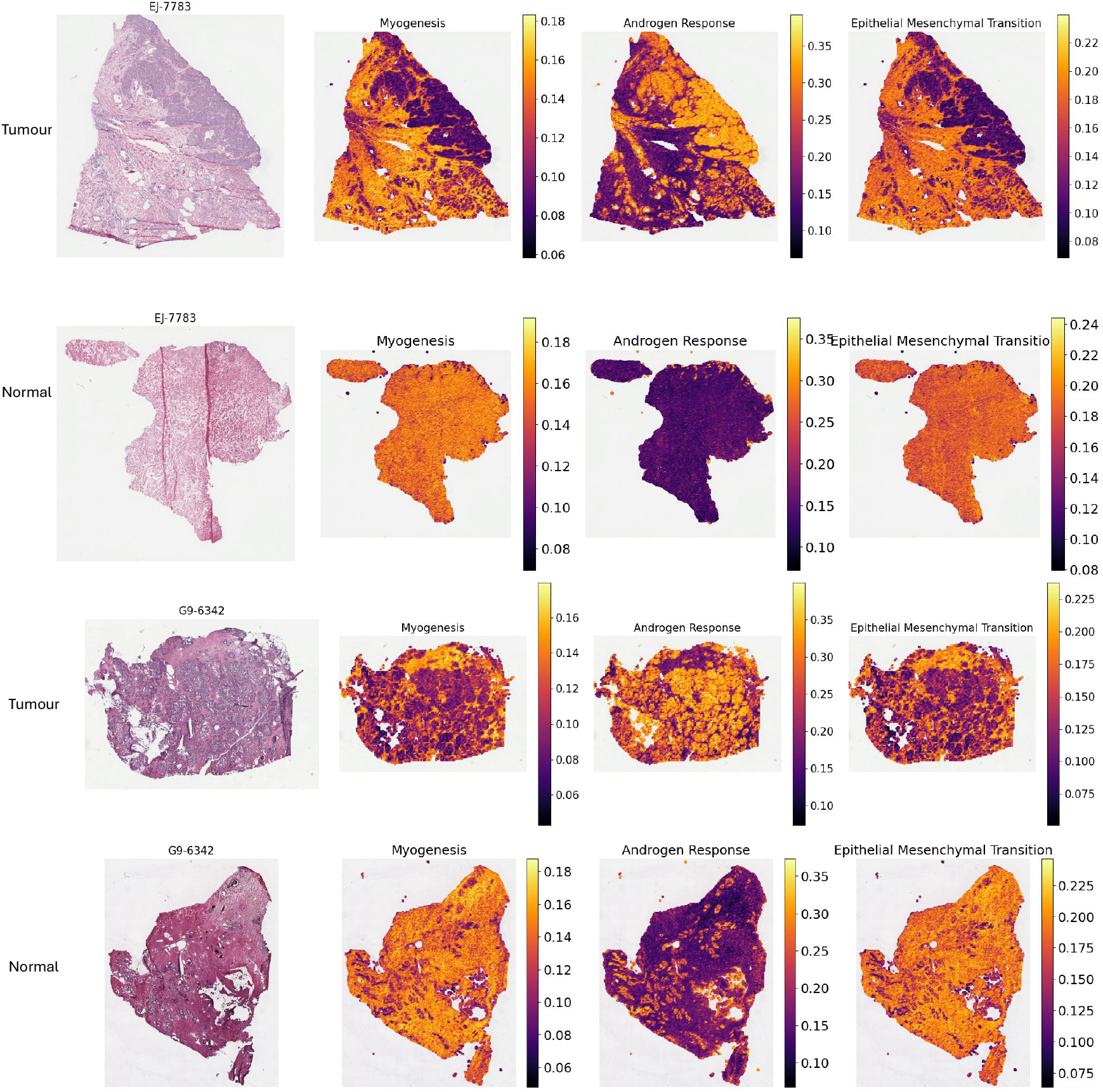
Spatial visualization of predictions obtained from the proposed model for Myogenesis, Androgen Response, and EMT pathways for TCGA patients having both tumour and normal tissue sections.

### D. Prediction of hypoxia signatures from H&Es of brain tumour patient samples

Hypoxia is one of the hallmarks of cancer and it is caused by the unbalanced oxygen availability due to insufficient blood supply and the excessive metabolic demand of tumour cells. Hypoxia causes substantial pressure on the tumour and its microenvironment leading to immune evasion and metabolic adaptation of neoplastic cells, maintenance of tumour cell stemness and resistance to treatment. It is also known to promote tumour progression and metastatic spread [45]– [48] Hypoxia can be traced in tissues using pimonidazole (PIMO) which binds selectively thiol-containing proteins or forms metabolites after reduction of its nitro group macromolecules when oxygen concentration falls below 10 mmHg [49].

Here we applied DeepPathway to samples from patients with GBM who received PIMO as part of a clinical trial [50]. Consecutive sections were stained for H&E and PIMO as ground truth and analysed with DeepPathway. We selected five brain tumour specific pathway definitions from Ravi et al. [34], which include (i) Reactive Hypoxia, (ii) Radial Glia, (iii) Reactive immune, (iv) Regional Neural Progenitor Cell (NPC), and (v) Regional Oligodendrocyte Precursor cell (OPC). Additional two pathway definitions we included are the subset of Reactive hypoxia definition, where one definition include 20 well established hypoxia specific genes while other subset include 41 genes (sum of 20 well established and 21 genes associated with hypoxic genes), curated locally. We trained DeepPathway model on five GBM samples (Ravi et al. [34]). After training the stage-1 model, we used the same GBM samples to train the stage-2 model. We used in-house brain-tumour patients H&Es to predict the hypoxia pathway expression. Results for three GBM patients having hypoxia clearly marked with PIMO stain are shown in Fig. 6 which shows that DeepPathway is able to mark certain hypoxic areas using only H&Es which is typically much harder task for the pathologist (See Supplementary Fig. 17 for other two patients). This approach provided a robust validation of DeepPathway and of its translational potential, providing a good visual concordance with the PIMO stain despite being trained on limited and low-quality ST brain tumour specific data. Indeed this will need further validation for model generalisation on routine H&Es going forward for any translational applications.

**Fig. 6:**
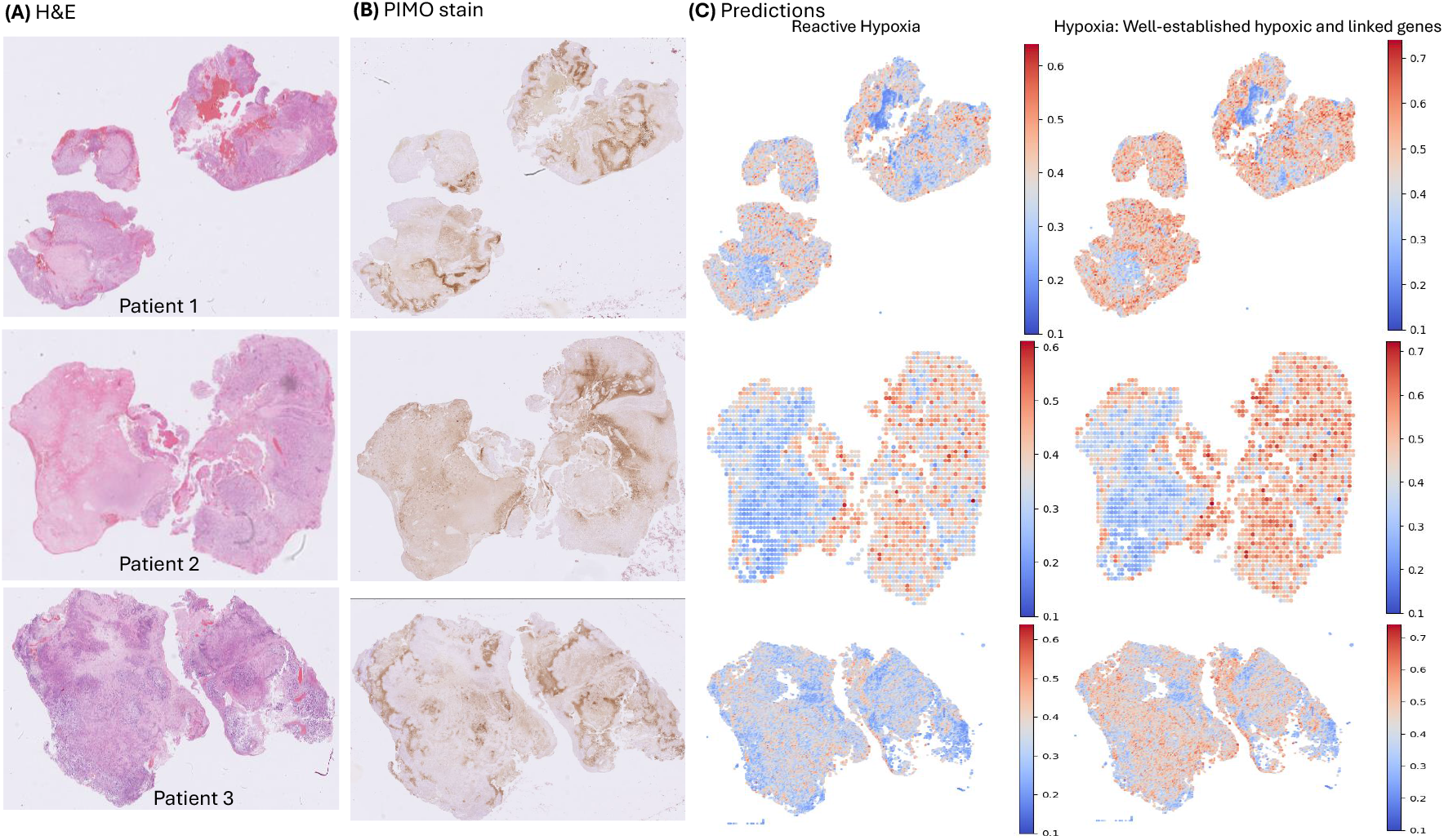
**(A)** Visualization of H&E image, **(B)** PIMO staining validation, and **(C)** Predicted Reactive Hypoxia pathway and a custom hypoxia geneset comprised of well-established hypoxic genes along with their linked genes from H&Es of brain tumour patients (row-wise) using DeepPathway.

## IV. Discussion and conclusion

In this paper, we proposed DeepPathway, a novel approach for the prediction of pathway expression from H&E images. Trained on ST data - this approach successfully links visual patterns in histological images to functional biological processes. We have applied it on publicly available cancer ST and imaging datasets, demonstrating its utility as a powerful tool for discovering how a cancer’s appearance relates to its underlying molecular state. H&E images are routinely available but their connection to functional biology remains largely uninterpreted. Existing AI models that predict expression of individual genes from histology fail to capture the cooperative nature of genes acting in pathways and can be noisy and biologically un-intuitive. DeepPathway addresses these gaps by linking tissue morphology and functional biology, aiming to extract a rich, pathway-level understanding of a tumour’s biology directly from a standard, low-cost image.

Our method builds on existing work for the prediction of gene expression from H&E images, utilising ST data (gene expression with co-registered images). We establish the utility of pathway prediction directly from H&Es, as compared to individual gene expression prediction followed by pathway quantification. For both gene and pathway expression prediction tasks, quality of images and their pre-processing could have significant impact on the model development and performance. DL-based expression prediction models mainly use spot-level H&E image patches, which are cropped around the center of the spots. However, in ST data, tissue sections may have variable image resolution due to the technical differences between experiments or variable spot diameters even using the same technology like 10X Visium. This leads to H&E patch sizes with inconsistent morphological features, i.e., cell shapes, density, and their surroundings. Additionally, in clinical settings, tissue H&Es without ST data can be stored in standard imaging formats like portable network graphics (PNG), where the image resolution in microns per pixel (MPP) is unavailable. To achieve the constant-sized image patch which is not affected by the spot diameter or the type of image, we have used TIA toolbox which provides patches of same size (say 224 × 224) with any given MPP resolution (say 0.5). Additionally, large-scale ST datasets, e.g. HEST-1k provides normalized and transformed WSIs in a pyramidal objects (TIFF) along with the metadata. However, in future, a multi-resolution DL-based framework for leveraging different magnifications will be needed to achieve more generalizable results by incorporating the images at any resolution.

Besides images, pre-processing and batch correction of multi-sample gene expression data might be needed before training the DL models for gene or pathway expression prediction. There are some statistical methods like Harmony [51] or probabilistic methods like single-cell variational inference [52] that uses the autoencoders based approach to remove batch effects. These methods transforms the gene counts data into another embeddings space which may contain negative gene expression values. However, in our work, we have not applied any batch correction method for the sake of obtaining positive valued gene expression prediction that are used to obtain reconstructed pathway expression. We also believe that transforming the gene expression data into pathway expression data using UCell reduces the batch effects because UCell is a rank-based method, which can handle the global shifts in gene expression levels caused by batch effects.

Recently, H&E foundation models are being widely applied to extract rich histology features as they are trained on enormous amount of WSIs. We have used the first version of optimus foundation model, i.e., *H-optimus-0*, however, there are other foundation models [53], [54] along with *H-optimus-1*^2^ (the model is trained on histology images sampled from over 1 million slides of more than 800, 000 patients) can be used as a future work. These models could lead to development of Za pathway expression foundation model that can be used on different tissues, i.e., brain, liver, or lungs, etc.

There exists patient-level heterogeneity in data which needs to be considered while developing an AI-based models for healthcare applications. For DeepPathway, we conducted experiments using single patient data samples and mixed samples that are taken from two different patients which shows that there is difference in performance when the model is trained on combined heterogeneous data rather than patient specific data. However, when we tested only on samples obtained from patient 1 (P1), we observed relatively low performance in terms of accurately predicting pathways. On the contrary, patient 2 (P2) samples achieved consistent performance when used under leave-one-out settings or in mixed patient samples approach. Upon further investigation, we can establish that low performance in P1 could be due to either very little or no spatial patterns in pathway expression which will make it harder for the contrastive learning model to learn localised patterns. Or, different H&E image resolutions between patient samples, e.g., 20X or 40X will give different patch view of the input data, which will affect model training. Furthermore, although both patients were diagnosed with Gleason 4+3 scores, however, there is a significant age difference in both patients, i.e., P1 is 82 years and P2 63 years old [32].

Furthermore, we show the utility of our model on TCGA data where there are only H&E images available besides bulk RNA data. Our method can show spatially localized expression of different pathways using H&Es, differentiating expression on tumour vs normal tissue slides for same patients. This can add further insight into the analysis of those samples, and could be explored further in conjunction with the bulk RNA data and other metadata available with those samples as a future application.

Lastly, we focused on GBM samples as a paradigm of hypoxic tumour. Any implementation of computational methods to identify hypoxia in H&E image will improve the ability to stratify patients at risk of a worse outcome. Noteworthy, hypoxia can only be identified using the stain for HIF-1*α* as a surrogate biomarker of low oxygen concentrations. At the same time, this was an excellent opportunity for validation and potential clinical utility of our method, as we have access to these precious human brain tumour samples which were collected in a way that they could be stained with PIMO as a ground truth for hypoxia. Our method makes good visual concordance with the PIMO stain despite being trained on limited and low-quality ST brain tumour specific data. These analyses can be further extended and validated to make the case for potential clinical applications for prediction of hypoxia in many cancer tissues.

Overall, to the best of our knowledge, this is the first approach to predict pathways from H&E images, and we believe with more training data, better quality images, refined pathway definitions, it will lead to further improvement and wider translational impact. Furthermore, we will extend DeepPathway for high-definition ST platforms such as Visium HD, which offers near-cellular resolution as future work.

## Acknowledgment

Muhammad Ahtazaz Ahsan was funded by MRC DTP award (MR/W007428/1).

We would like to thank all authors of the public data that is used in this study. We also thank Gabriella Forte for her support in the preparation of the in-house brain tumour histology slides and PIMO staining data.

## Footnotes

1 https://www.genome.gov/about-genomics/fact-sheets/Biological-Pathways-Fact-Sheet

2 https://huggingface.co/bioptimus/H-optimus-0-1

