## Supplementary figures for "DeepPathway: Predicting Pathway Expression from Histopathology Images"

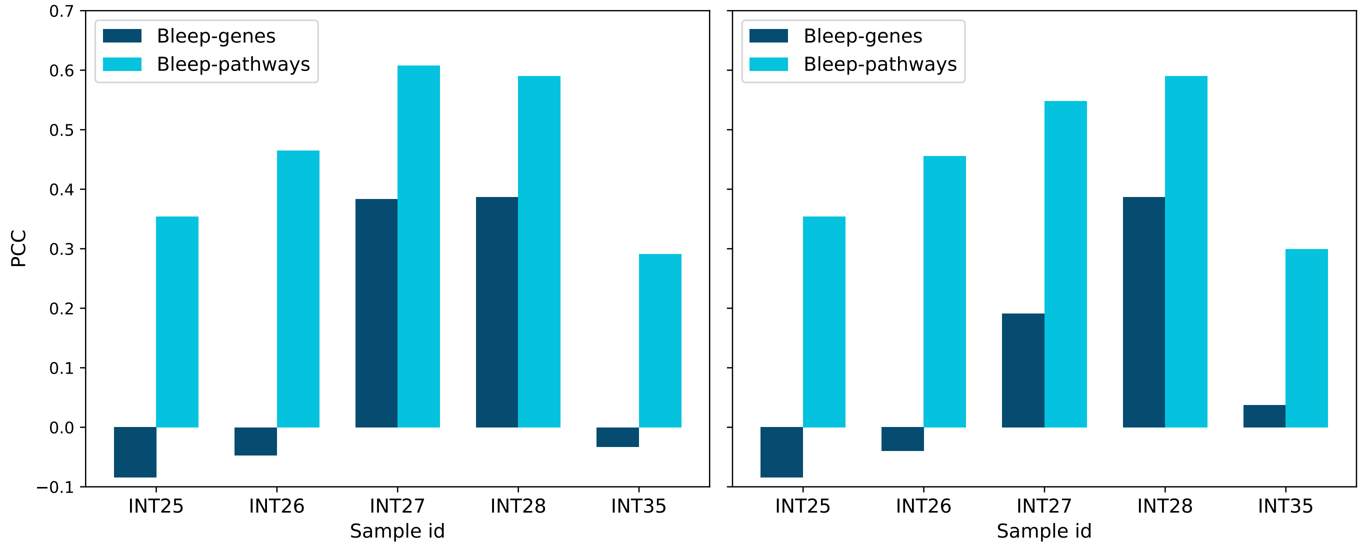


Fig. 1: Average PCC of top 5 highly expressed **(left)** and highly variable **(right)** pathways obtained from baseline method, i.e., BLEEP-pathways and BLEEP-genes, on Small prostate data.

* Joint corresponding authors

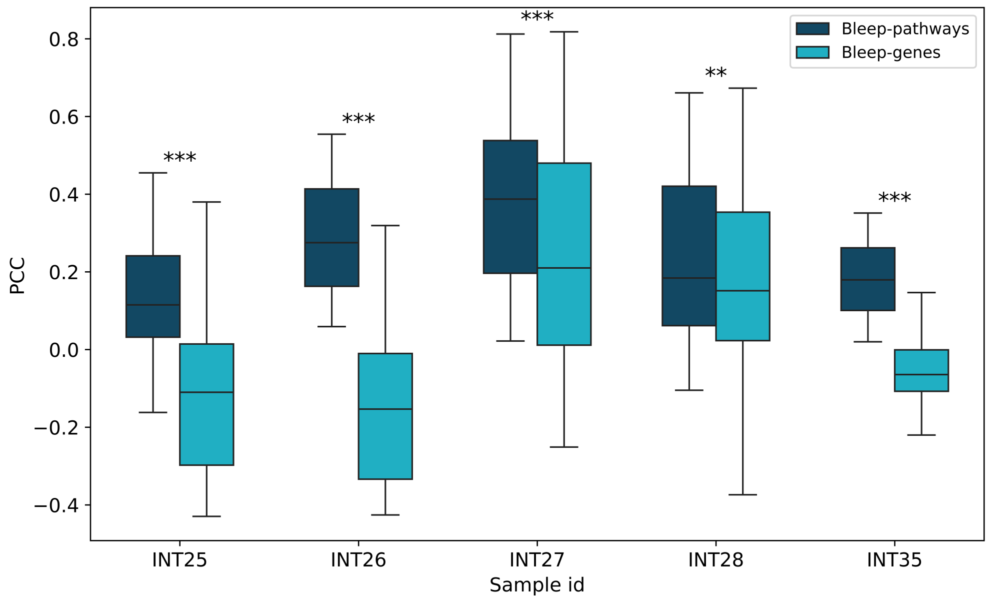


Fig. 2: PCC distribution of Bleep-pathways model when trained with ResNet18 as an image encoder for small prostate cancer dataset.

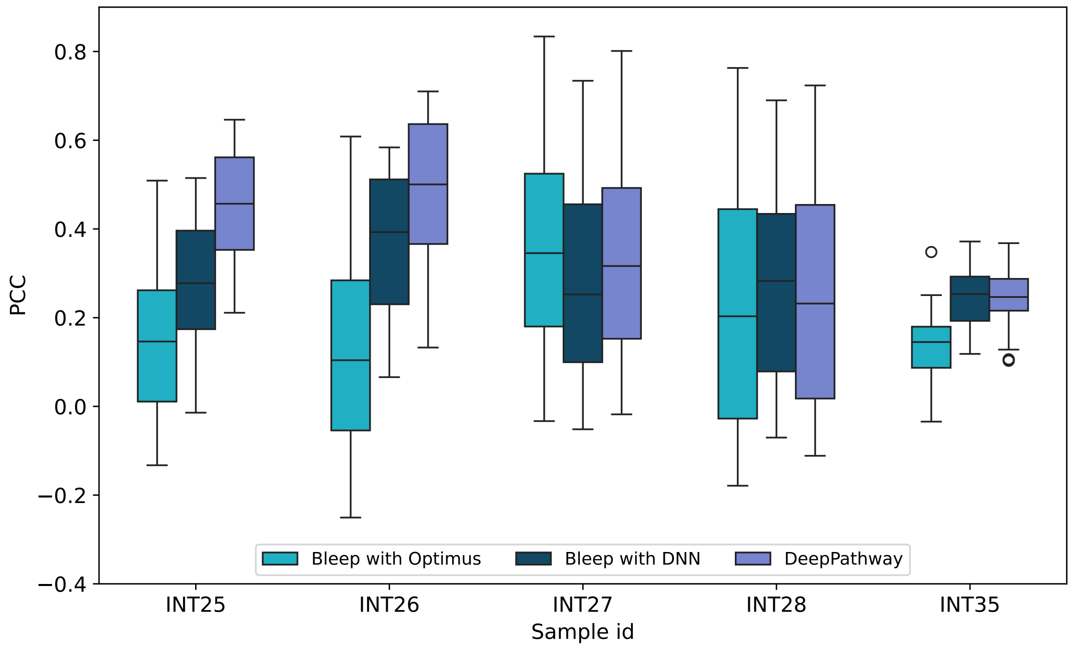


Fig. 3: Validation study of DeepPathway (BLEEP with Optimus and DNN) by removing different components of the model on small prostate cancer data where image encoder is ResNet18 for every model.

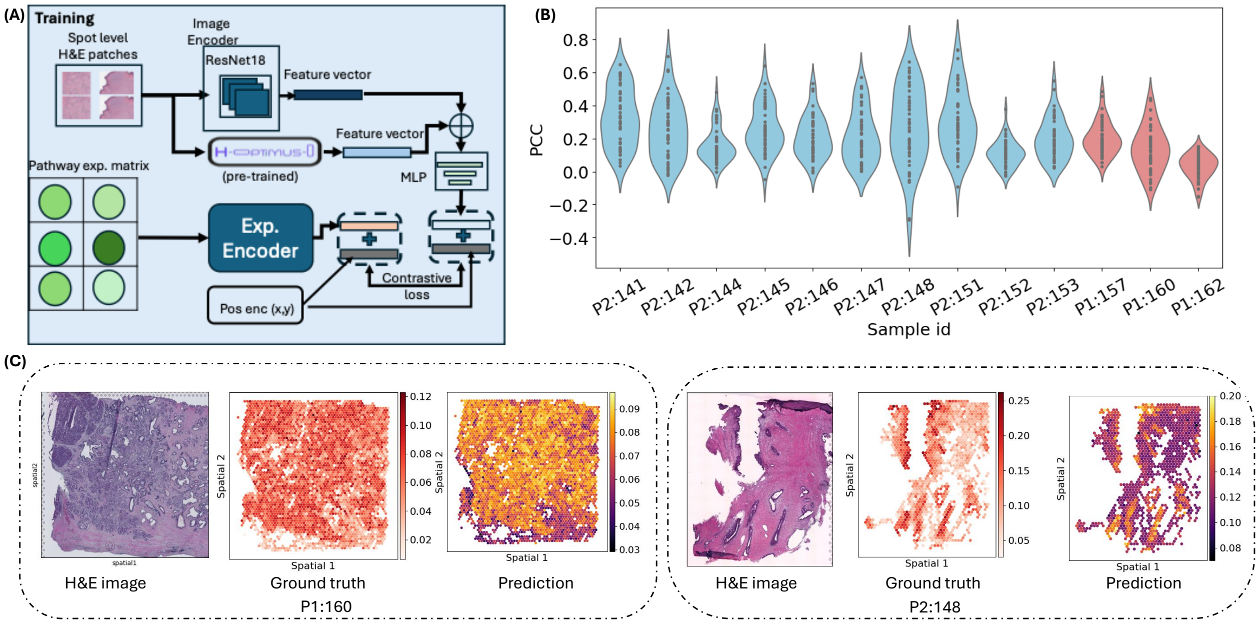


Fig. 4: **(A)** Integration of H-Optimus-0 H&E foundation model with BLEEP (Stage-1 of DeepPathway model) to predict pathway expression from mixed samples of Patient 1 and Patient 2 of large prostate cohort. A combined 2048-dimensional feature vector (1536 from non-trainable H-Optimus-0 foundation model and 512-dimensional from trainable ResNet18) is inputted to BLEEP for model training and validation. **(B)** PCC distribution plot of test samples taken from Patient 1 and Patient 2 **(C)** test H&E image and spatial visualization of the heme Metabolism (P1:160) and p53 pathway (P2:148) ground truth pathway expression—obtained from UCell—alongside the predicted pathway expression. The predicted expression achieved a PCC value of 0.44 and 0.67 for, respectively.

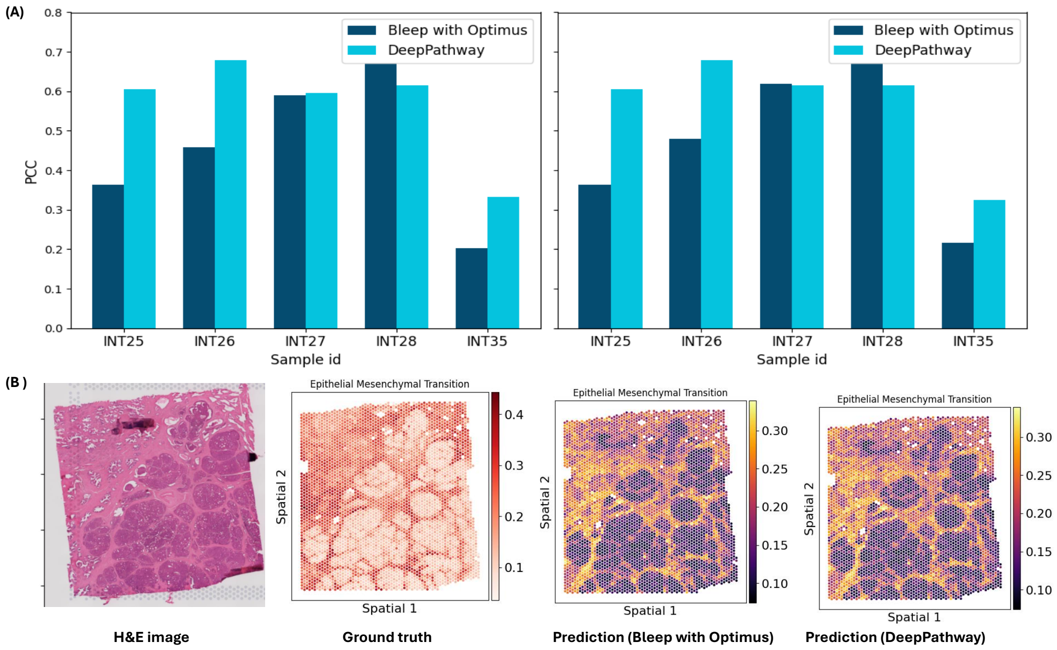


Fig. 5: **(A)** Average PCC of top 5 highly expressed **(left)** and highly variable **(right)** pathways obtained from Bleep with Optimus (Stage-1) and DeepPathway. **(B)** H&E and spatial visualization of the ground truth (obtained from the UCell) and predictions obtained from both models. The predicted expression achieved the PCC of 0.83 and 0.80 for the Epithelial Mesenchymal Transition pathway (sample ID=INT27).

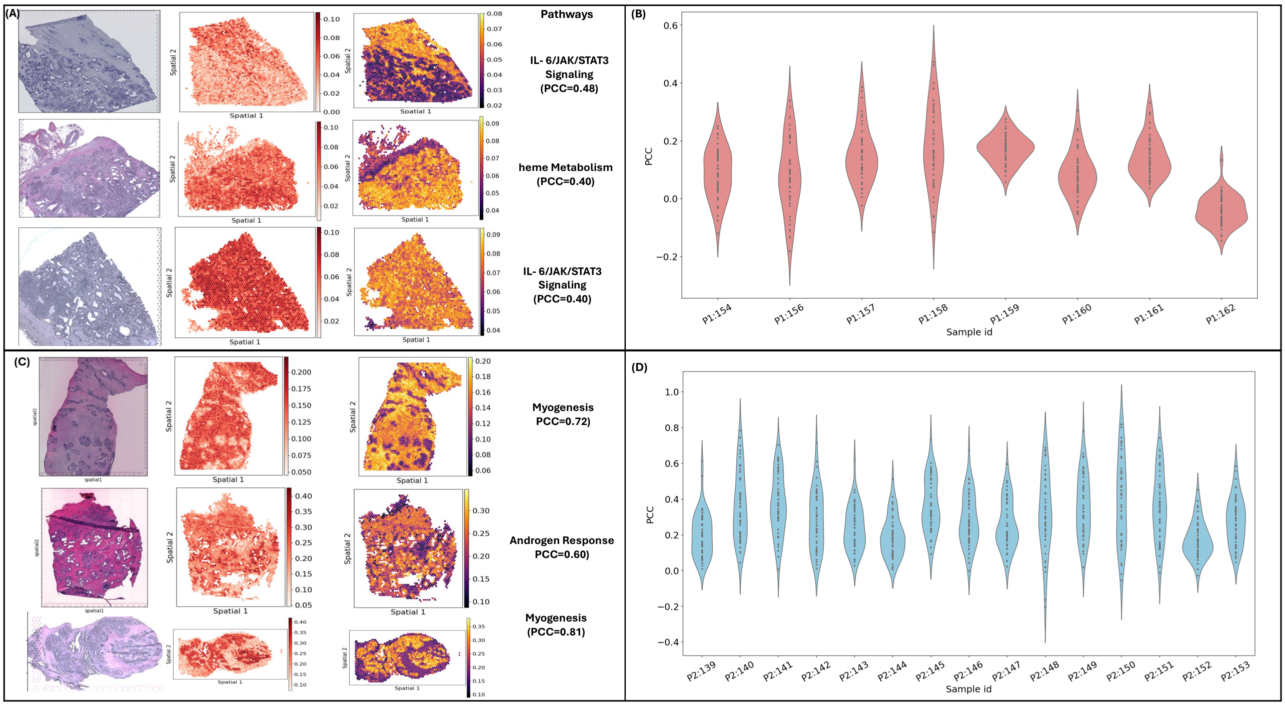


Fig. 6: Patient-wise evaluation of Bleep with Optimus (Stage-1 of DeepPathway model) using leave-one-out cross validation. **(A)** H&E along with spatial visualization of ground truth and prediction obtained for best predicted pathways from Patient 1 samples (Ids=157,158,161). **(B)** PCC distribution of all pathways using leave-one-out cross validation for Patient 1. **(C)** H&E along with spatial visualization of ground truth and prediction obtained for best predicted pathways from Patient 2 samples (Ids=142,147,150). **(D)** PCC distribution of all pathways using leave-one-out cross validation for Patient 2.

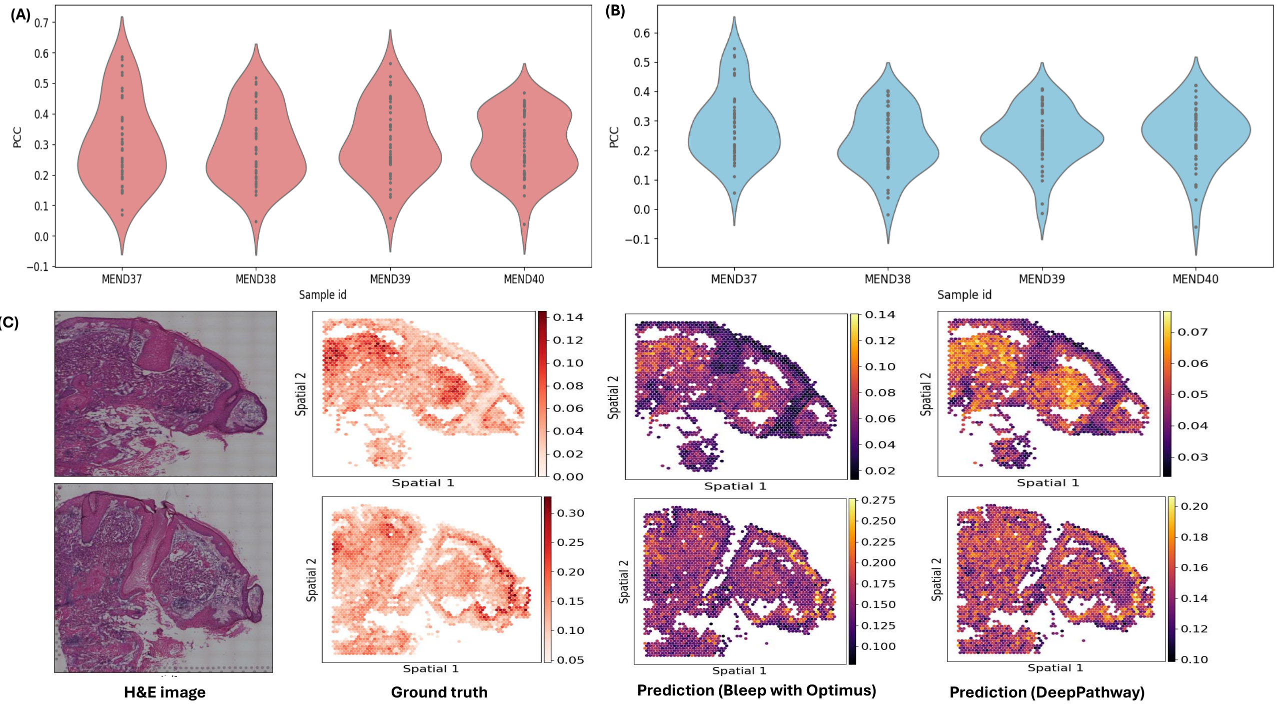


Fig. 7: **(A)** PCC distribution of each sample in leave-one-out cross validation on skin cancer data using Stage-1 of DeepPathway (BLEEP with *H-Optimus-0* model). **(B)** PCC distribution of each sample in leave-one-out cross validation on skin cancer data using DeepPathway **(C)** H&E and spatial visualization of the ground truth (obtained from the UCell) and predictions obtained from both models. The predicted expression achieved the PCC of 0.59 and 0.52 for stage-1 and stag-2, respectively, for E2F Targets pathway (top row, sample ID=MEND37), while achieved the PCC of 0.52 and 0.41, for Myc Targets V1 pathway (bottom row, sample ID=MEND39).

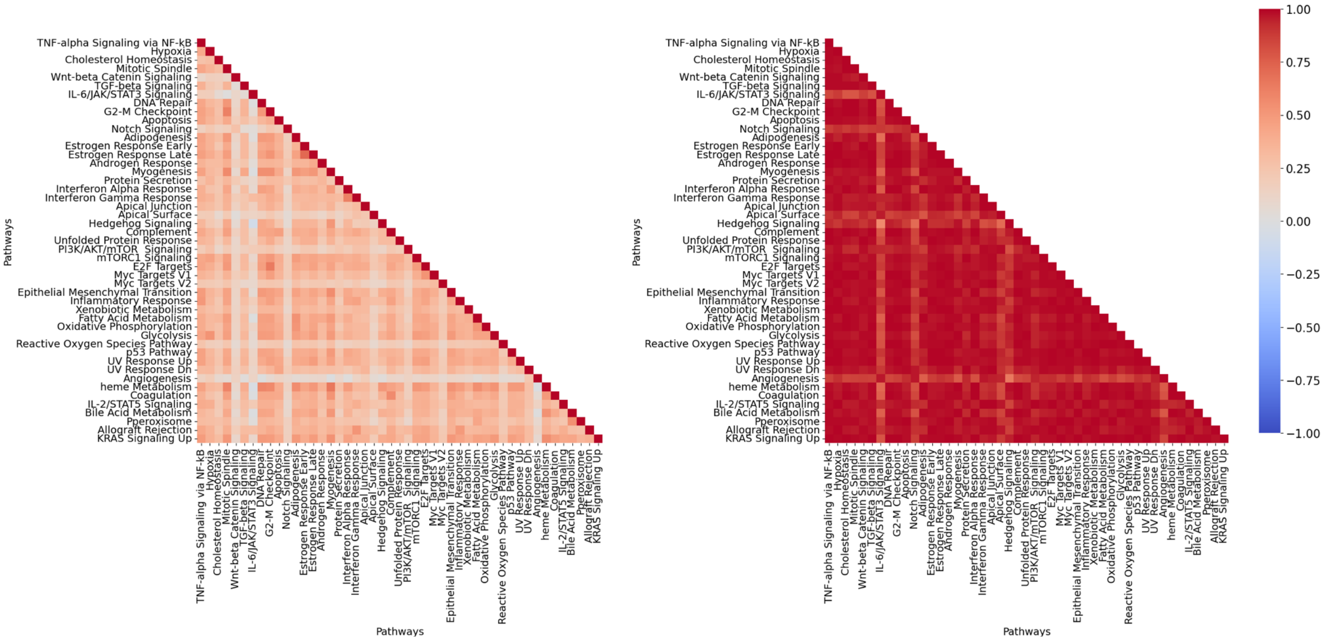


Fig. 8: intra-correlation computed between ground truth and predicted pathway expression (from DeepPathway model trained on Patient 1 samples only) for sample id=158, computed using 47 pathway definitions obtained from MSigDB hallmark on large prostate cancer data.

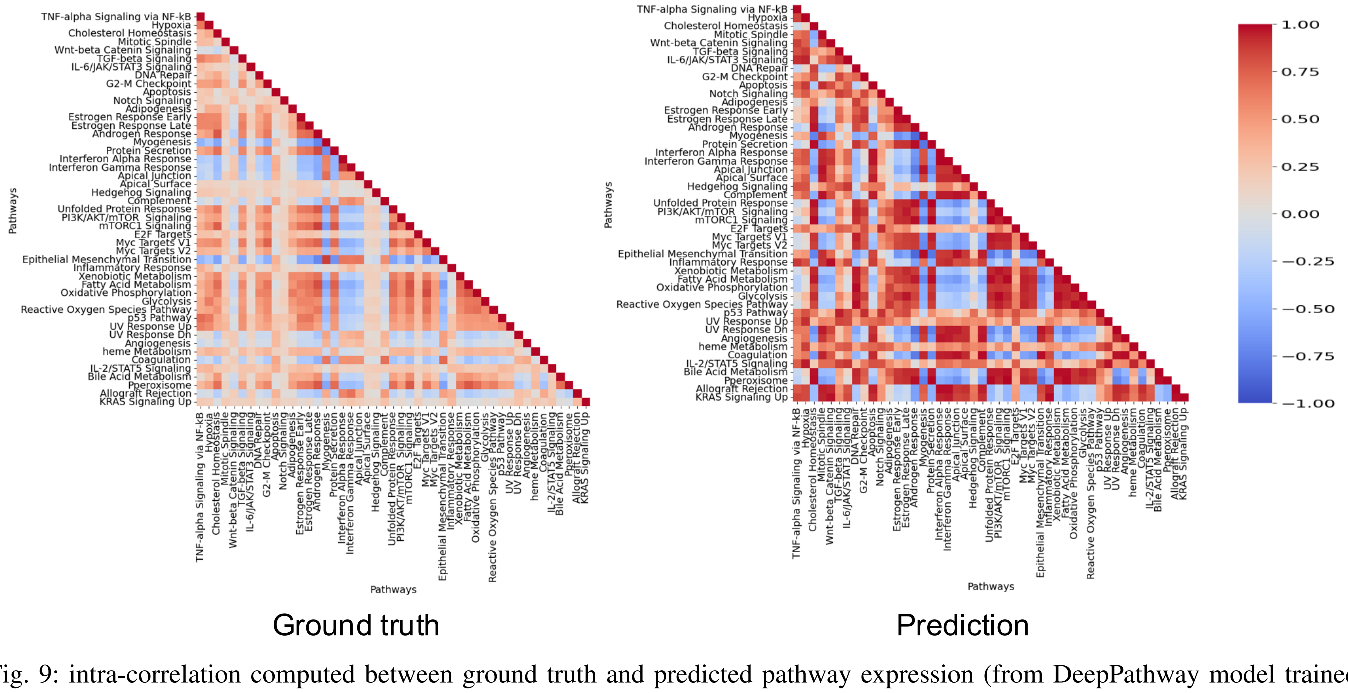


Fig. 9: intra-correlation computed between ground truth and predicted pathway expression (from DeepPathway model trained on Patient 2 samples only) for sample id=150, computed using 47 pathway definitions obtained from MSigDB hallmark on large prostate cancer data.

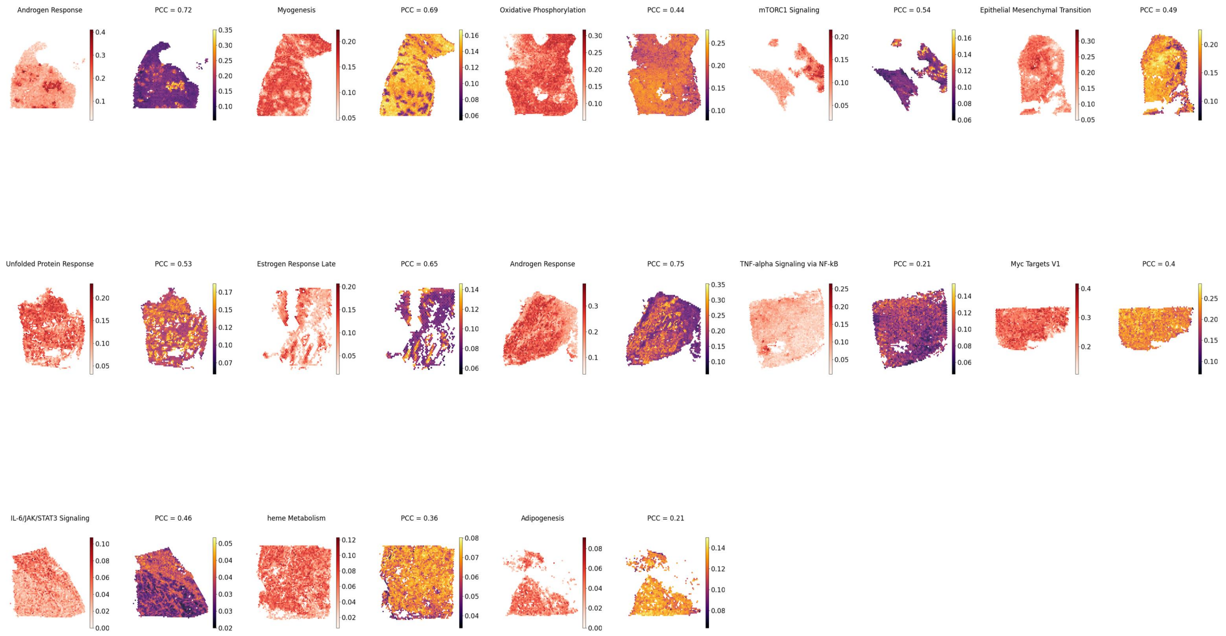


Fig. 10: Spatial visualization of pathways predicted well within all slides (selected from top 5 highly predicted pathways) from DeepPathway model using mixed samples from Patient 1 and Patient 2.

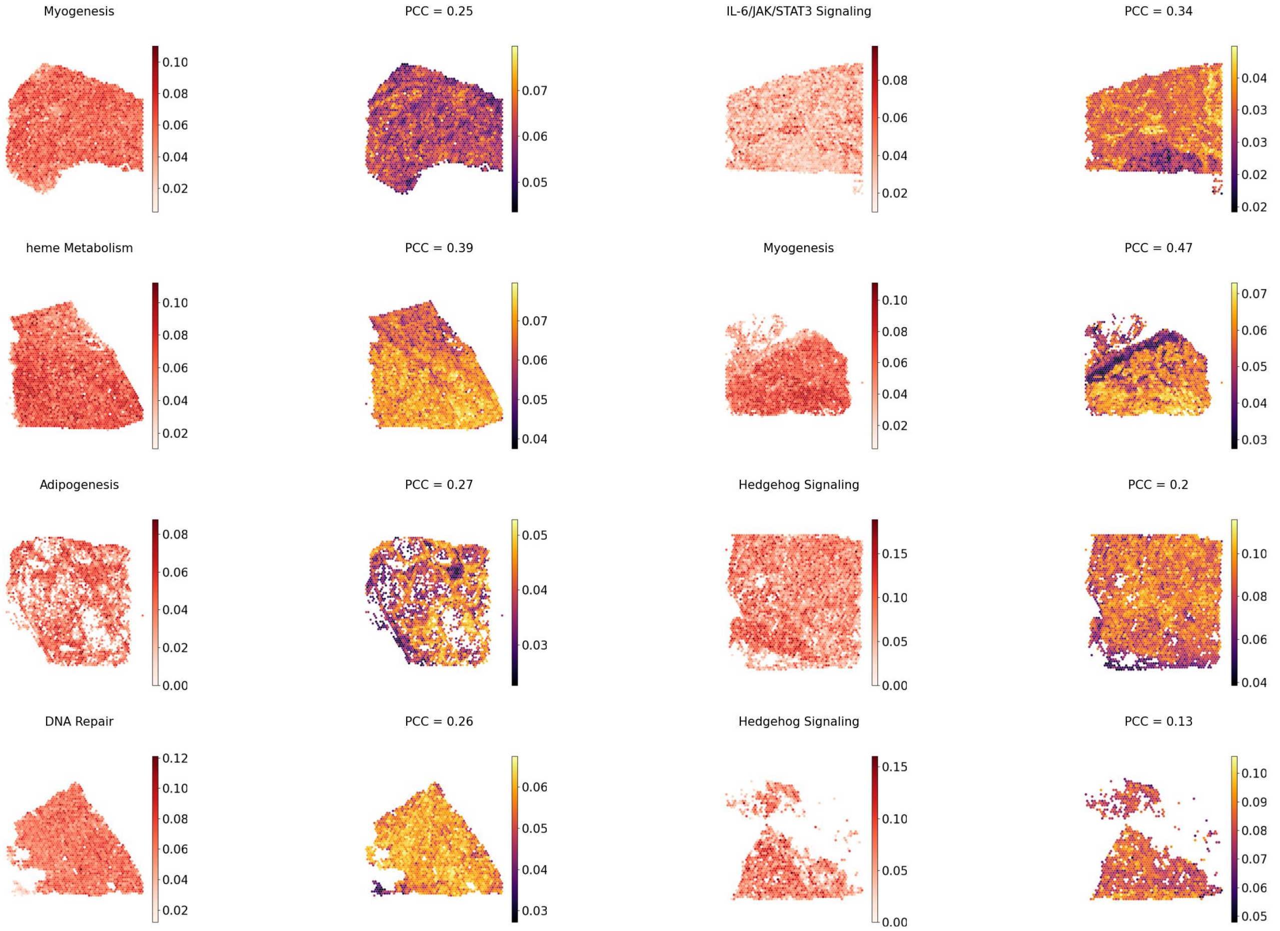


Fig. 11: Spatial visualization of pathways predicted well within all slides (selected from top 5 highly predicted pathways) from DeepPathway model using large prostate cancer data Patient 1 only with leave-one-out cross validation.

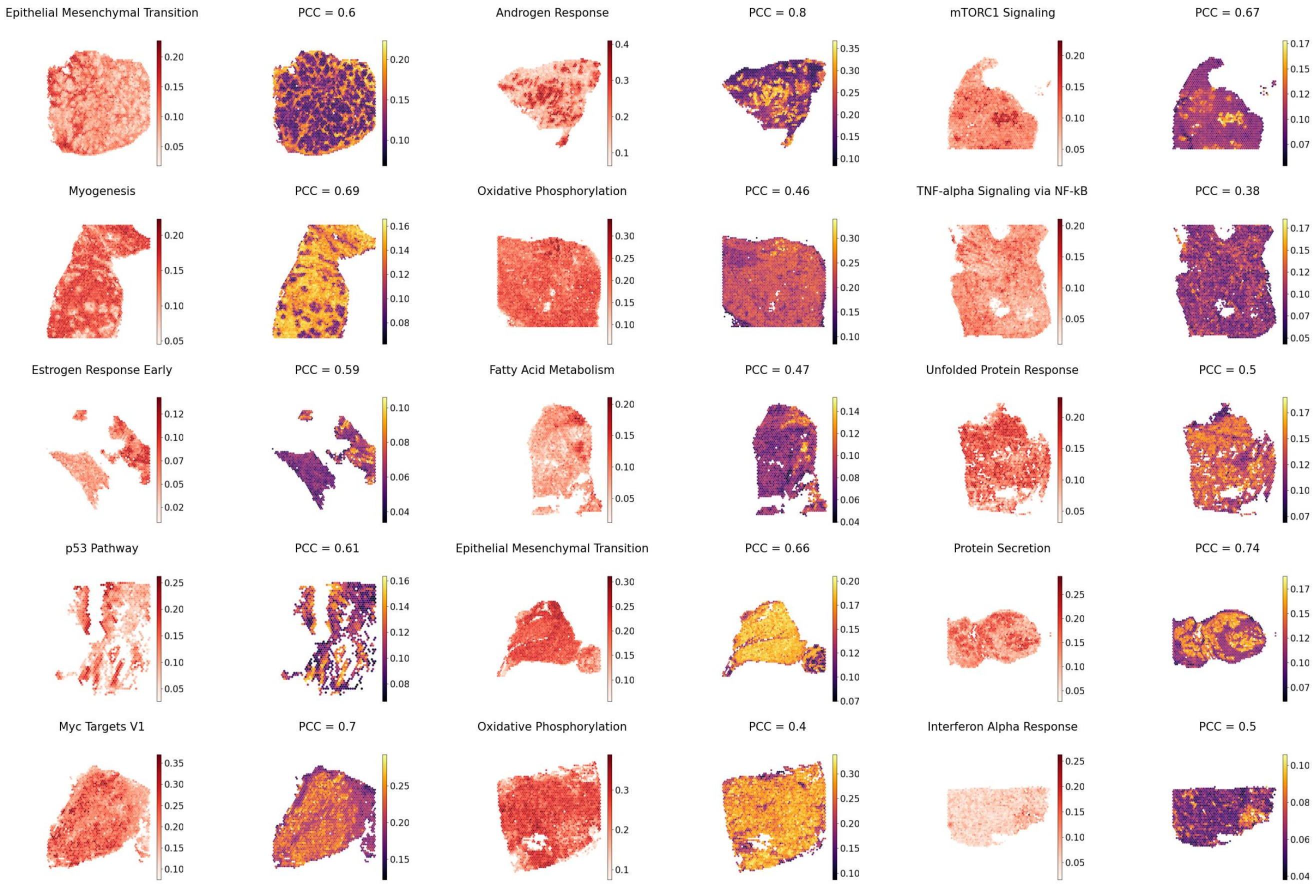


Fig. 12: Spatial visualization of pathways predicted well within all slides (selected from top 5 highly predicted pathways) from DeepPathway model using large prostate cancer data Patient 2 only with leave-one-out cross validation.

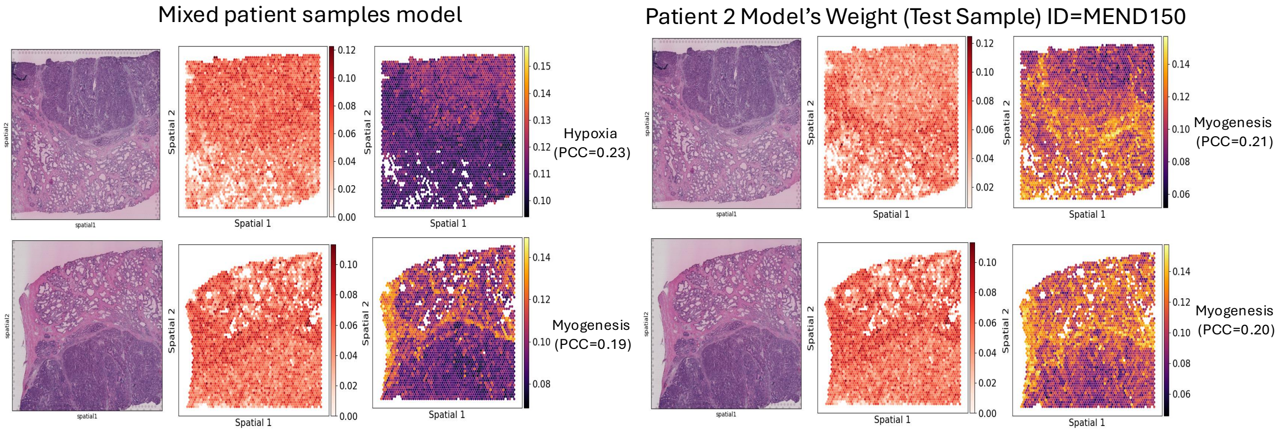


Fig. 13: Evaluation of DeepPathway model using two samples, i.e., MEND61 (top-row) adn MEND62 (bottom-row) from another prostate cancer study (taken from HEST-1k) data as an independent data cohort where the model is trained (i) using mixed patient samples of P1 and P2 **(left)** and (ii) using model trained only on samples of P2 (**(right)**) for showing its robustness and generalizability.

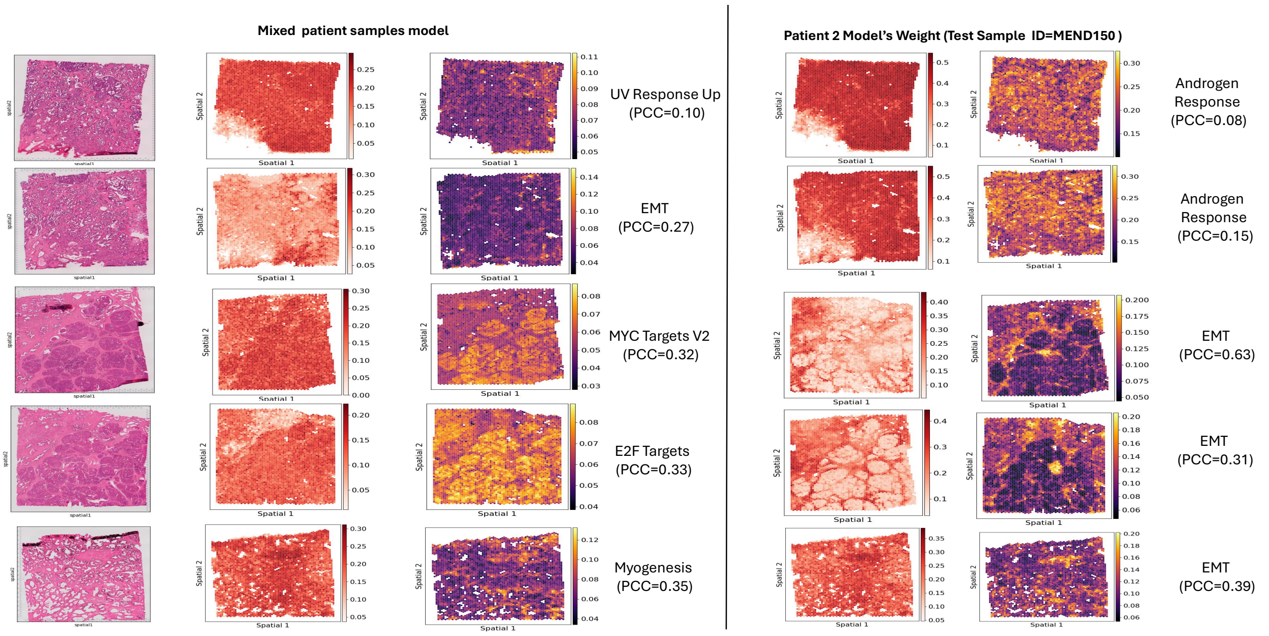


Fig. 14: Evaluation of DeepPathway model using small prostate data as an independent data cohort where the model is trained (i) using mixed patient samples of P1 and P2 9**(left)** and (ii) using model trained only on samples of P2 (**(right)**) for showing its robustness and generalizability.

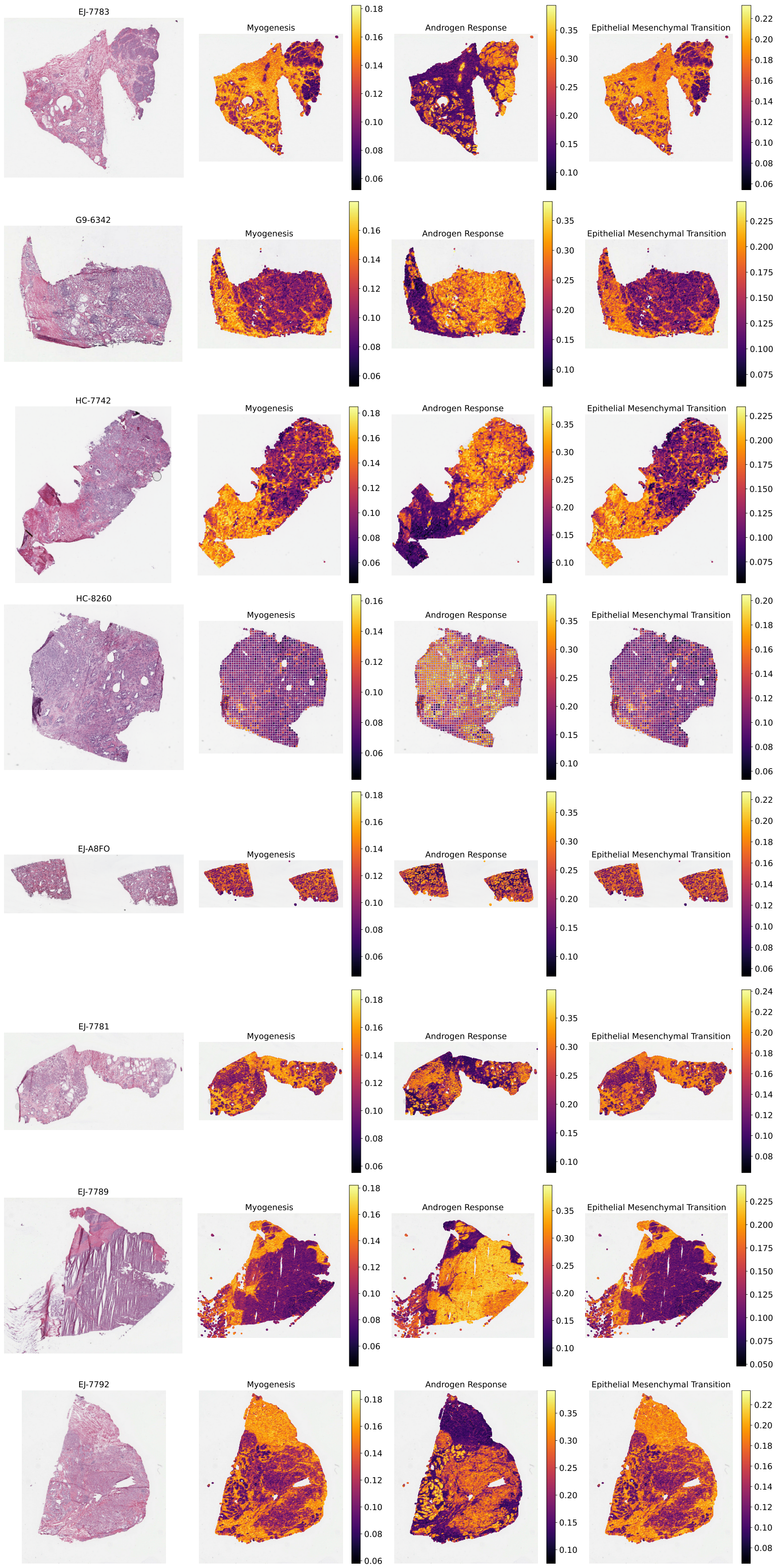


Fig. 15: Visualization of Tumour TCGA prostate H&E slides.

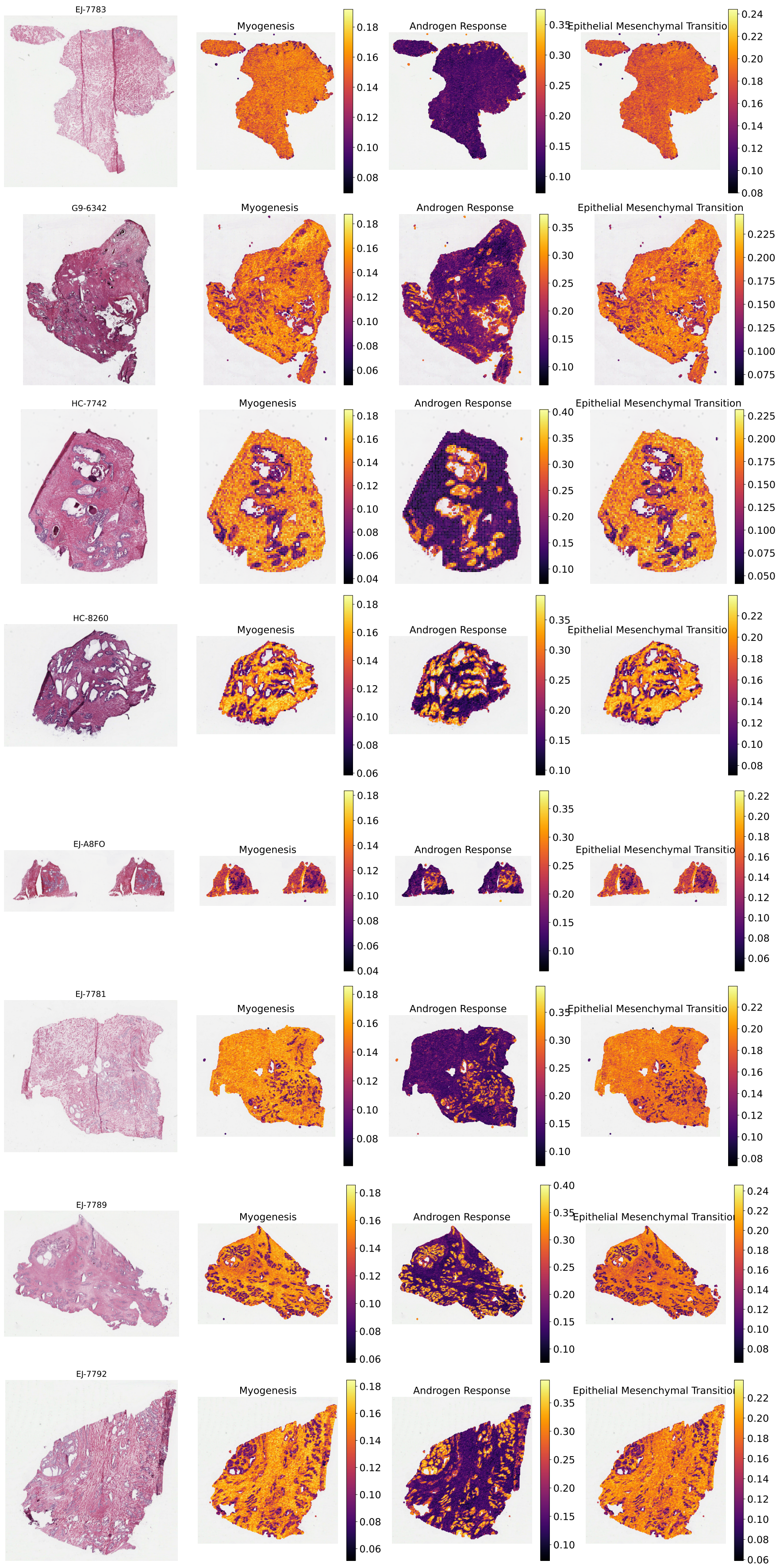


Fig. 16: Visualization of Normal TCGA prostate H&E slides.

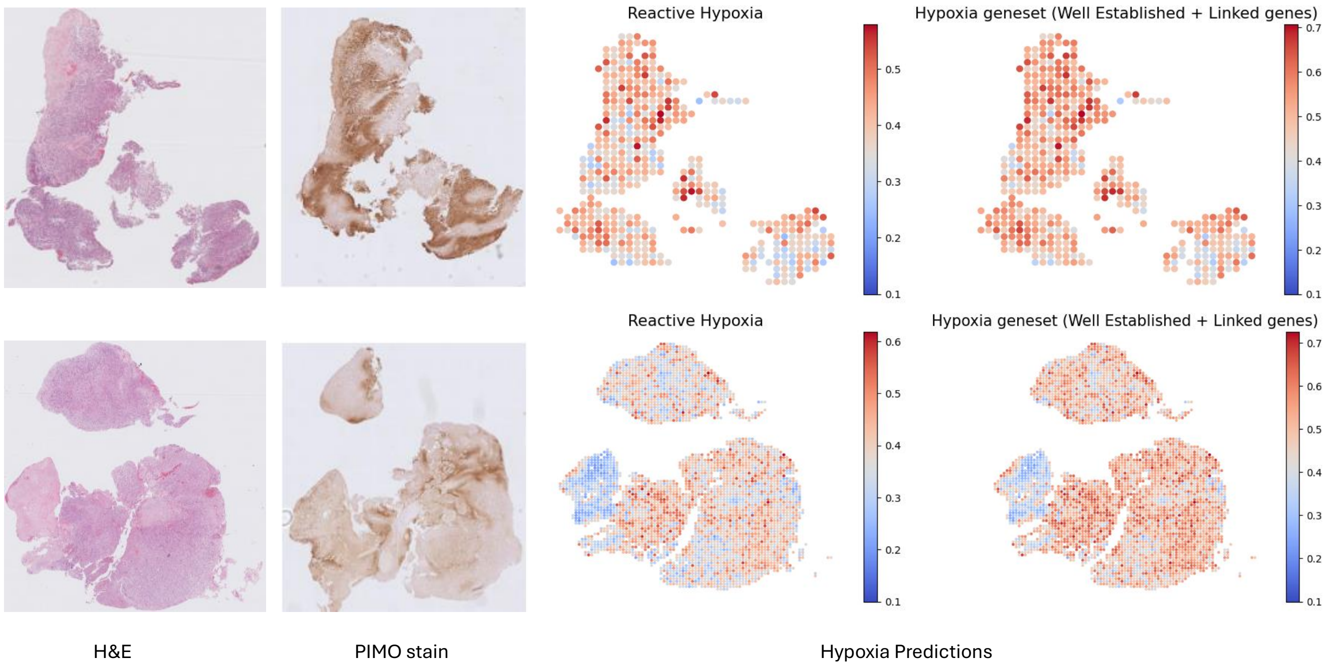


Fig. 17: Visualization of H&E images, PIMO stainin g validation, and predicted Reactive Hypoxia pathway and a custom hypoxia geneset comprised of well-established hypoxic genes along with their linked genes from H&Es of brain tumour patients (row-wise) using DeepPathway.
